# Leveraging the salivary microbiome profile to stratify REM sleep behavior disorder and synucleinopathies

**DOI:** 10.1101/2025.10.26.684688

**Authors:** Daisuke Hisamatsu, Hiroaki Masuoka, Haruka Takeshige-Amano, Taku Hatano, Takashi Ogawa, Kazuo Yamashiro, Rina Kurokawa, Yo Mabuchi, Yuna Naraoka, Takashi Asada, Wataru Suda, Chihiro Akazawa, Nobutaka Hattori

## Abstract

Few studies have explored noninvasive biomarkers of synucleinopathies, including Parkinson’s disease (PD), and rapid-eye-movement behavior disorder (RBD), a prodromal stage for these conditions. Human oral/salivary microbiomes are altered in PD, highlighting their potential role in both PD pathogenesis and diagnosis. We analyzed 249 salivary microbiomes of controls and patients with idiopathic RBD and various synucleinopathies, including two subgroups divided based on the presence of RBD symptoms, through both cross-sectional and longitudinal studies. The microbiome composition was strikingly similar between patients with RBD and early PD exhibiting RBD symptoms. The area under the curve range for distinguishing RBD from controls and each synucleinopathy was 0.85–0.94. We further performed pseudotime trajectory analysis of the microbiome compositional space; populations with low diversity, enriched *Streptococcus*, and depleted *Neisseria* exhibited a brief pseudotime transition from RBD to early PD. Our findings suggest that early PD-like salivary dysbiosis manifests during RBD, allowing for the stratification of synucleinopathies through the innovative use of salivary microbiome profiles.

---

Abnormal α-synuclein aggregation is a hallmark of synucleinopathies, such as Parkinson’s disease (PD), dementia with Lewy bodies (DLB), and multiple system atrophy (MSA). The propagation theory stands as the foremost hypothesis illuminating the aggregation mechanism of pathological α-synuclein, offering profound insights into its complex behavior. Alongside this theory, two routes have been proposed as hypotheses: (1) body-first type wherein aggregation begins in the enteric or peripheral nervous system and propagates through the vagus nerve to the central nervous system (CNS) and (2) brain-first type wherein aggregation begins in the CNS^1,2^. Idiopathic (isolated) rapid-eye-movement sleep behavior disorder (iRBD) is proposed as a prodromal stage of body-first type^1,2^. Pathological α-synuclein is aggregated in several peripheral tissues, including the gut^3,4^, salivary glands^5^, oral mucosa^6^, and skin^7,8^. The α-synuclein seeds are also detected in biofluids, such as cerebrospinal fluid and blood^9,10^, which are perceived as biomarkers of synucleinopathies. Additionally, metabolites in biofluids have attracted attention as potential biomarkers for PD diagnosis^11^. However, the biomarkers that can stratify synucleinopathies, including RBD, remain largely unknown.

Increasing evidence suggests a relationship between environmental factors, such as the gut microbiome, and α-synuclein aggregation and its propagation from systemic neurons to the CNS^12–14^. Additionally, recent studies have reported that the gut microbiome differs between patients with RBD and PD^15^, as well as between two groups of patients with PD categorized by RBD symptom presence or absence^16^. Similarly, alterations in oral/salivary microbiomes have been observed in patients with PD^17–20^. In our previous study, gut and salivary microbiome alterations were associated with PD progression^21^. Notably, the discriminative ability between patients with PD and Alzheimer’s disease (AD) was higher in the saliva than in the gut^21^. Considering α-synuclein accumulation in the salivary glands and oral mucosa of patients with iRBD^6^, we hypothesized that the oral microbiome from the prodromal stage would change before synucleinopathy onset. However, few studies have investigated oral/salivary microbiomes in synucleinopathies, including RBD. Thus, we aimed to demonstrate our hypothesis by identifying differences in salivary microbiome alterations among patients with iRBD and synucleinopathies, as well as the possibility of early diagnosis by leveraging the salivary microbiome. In this study, we took a step further in categorizing synucleinopathies by examining the presence of RBD symptoms, anticipating that the salivary microbiome will play a crucial role, not only in facilitating early diagnosis but also in unraveling the distinctions between body-first and brain-first PD pathogenesis.

RBD to PD conversion often takes more than 10 years^2,22^, making longitudinal analyses challenging. In cell biology, trajectories and pseudotime analyses based on gene expression profiles at the single-cell level are employed to better understand cell-state transitions (e.g., developmental processes)^23^. These analyses reduce high-dimensional data, including massive cross-sectional datasets, into a low-dimensional space based on the similarity of expression profiles between samples, using dimension reduction methods such as principal component analysis (PCA)^24^. Recently, trajectory analyses have been performed using the human microbiome^25,26^. Tap et al. used trajectory analysis of gut microbiome data throughout the human lifespan and found distinct clusters in the microbiome compositional space^25^. These clusters were associated with a proportion of specific genera (e.g., *Bacteroides*-enriched population) related to host factors. However, this study was not designed to address pseudotime labeling during disease progression. Tsamir-Rimon et al. developed a computational framework to detect the manifold (low-dimensional roots) in the microbiome compositional space associated with disease progression using human vaginal microbiome samples from both cross-sectional and longitudinal studies^26^. Thus, to examine the microbial and host features during RBD-to-PD progression, we applied this framework to cross-sectional and longitudinal samples of patients with iRBD and synucleinopathies.

## Results

### Salivary microbiome is altered in RBD and synucleinopathies

Saliva samples were collected from 249 participants, including patients with iRBD and synucleinopathies or non-neurodegenerative control individuals, from primary and test cohorts in Japan (Fig. 1a). For the longitudinal analyses, we repeatedly collected 23 saliva samples from patients with iRBD and *de novo* patients (naïve patients without anti-PD drug use) with PD (hereinafter “*de novo*”). A total of 272 samples were subjected to bacterial 16S ribosomal RNA (rRNA) gene amplicon sequencing and multivariate analyses incorporating medication and oral health variables. The demographics of the primary cohort are shown in Table 1. Although significant age differences were observed among the *de novo*, PD-RBD, MSA, DLB, or RBD groups (*P*<0.04; Table 1), only disease significantly contributed to the salivary microbiome composition in a stepwise redundancy analysis (RDA) based on the weighted UniFrac distance (cumulative adjusted *R*^2^=4.9%, false discovery rate [*FDR*]=0.009; Fig. 1b). To evaluate disease differences, we performed the weighted UniFrac distance-based β-diversity (inter-individual dissimilarity considers both the phylogeny and relative abundance of microorganisms) and visualized microbiome composition changes with principal coordinate analysis (Fig. 1c). Significant differences in the RBD group were observed in the control, DLB, or MSA groups (*P*=0.009, 0.009, and 0.001, respectively; Fig. 1d). Significant differences in patients with MSA were observed in the DLB and *de novo* groups (*P*=0.002 and 0.003, respectively; Fig. 1e,f). However, no significant differences in either synucleinopathy group were found between those divided based on the RBD symptom presence (Fig. 1g). To further explore the effect of RBD symptoms on the microbiome composition, we compared weighted UniFrac distances between the RBD and each group. The distances from RBD to synucleinopathy groups with RBD symptoms (*de novo*-RBD, PD-RBD, and MSA-RBD) were closer than those to the groups without RBD symptoms (*de novo*, PD, and MSA; Fig. 1h). The *de novo*-RBD group showed significant differences from the other disease groups, except the PD-RBD group (*P*<0.021; Fig. 1h). These results suggest that the salivary microbiome composition in patients with iRBD is similar to that of patients with PD having RBD symptoms than that of controls.

**Fig. 1.**
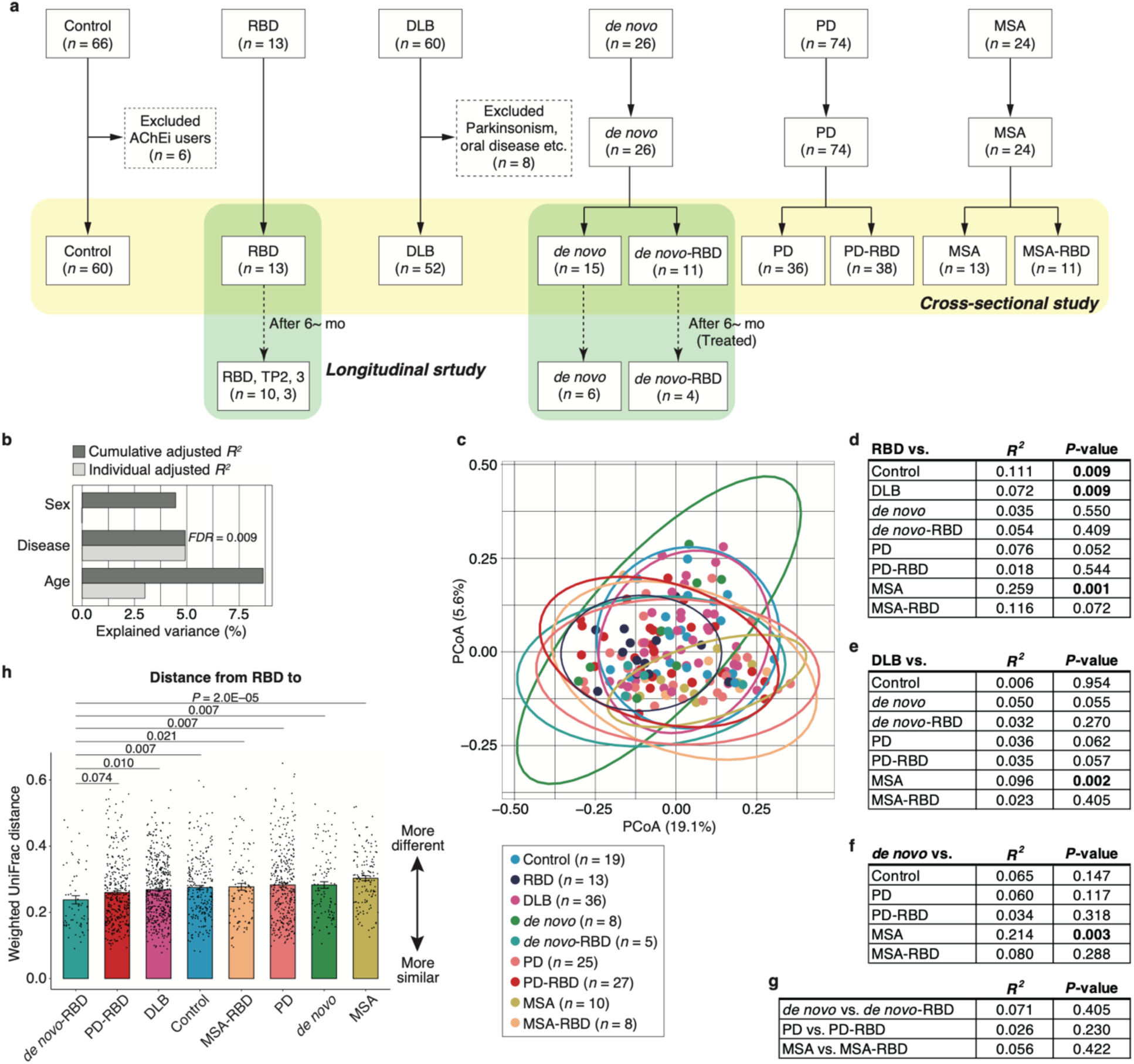
Microbial structure in patients with RBD is similar to that of patients with synucleinopathies with RBD symptoms. **a**, Flow chart of participant selection. **b**, Individual and cumulative adjusted *R*^2^ (explained variance) of variables (age, sex, and disease) in stepwise redundancy analysis based on the weighted UniFrac distance in the primary cohort (*n*=151). **c**, Graph showing weighted UniFrac- PCoA in the disease category. **d–g**, Permutational multivariate analysis of variance based on weighted UniFrac distance comparing each disease group. The numbers of participants are shown in the figure. The *R*^2^ and *P*-values were determined by permutational multivariate analysis of variance using the Benjamini–Hochberg method. **h**, Graph showing the mean distance calculated using the weighted UniFrac distance matrix between the RBD and each group. The dots indicate all combinations of participants included in the group. Statistical significance was determined using the Wilcoxon rank-sum test with the Benjamini–Hochberg method (*P*<0.05). AChEi, acetylcholinesterase inhibitor; *de novo*, *de novo* patients with PD; DLB, dementia with Lewy bodies; FDR, false discovery rate; MSA, multiple system atrophy; PCoA, principal coordinate analysis; PD, Parkinson’s disease; RBD, rapid-eye-movement sleep behavior disorder; TP, time point.

**Table 1.**
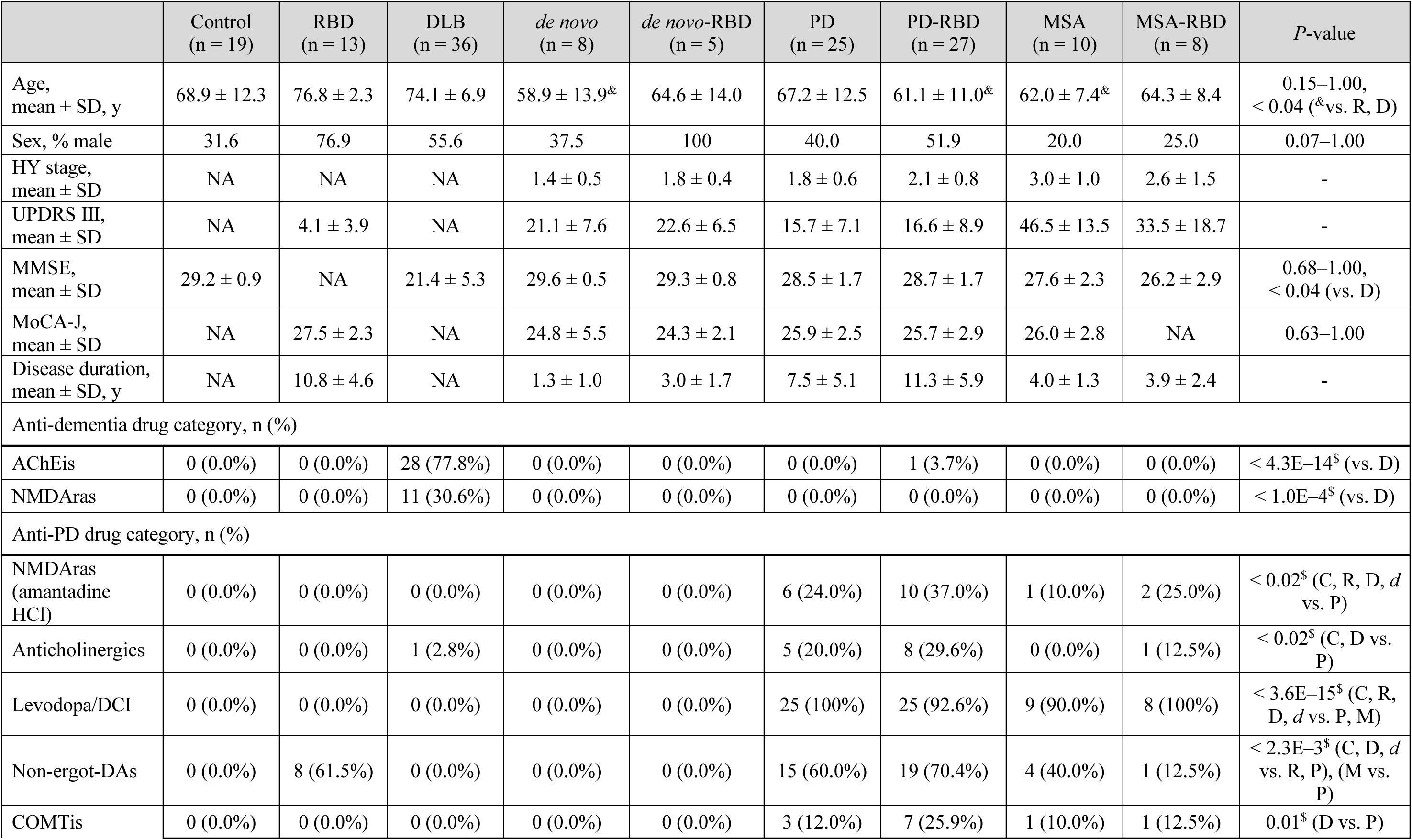
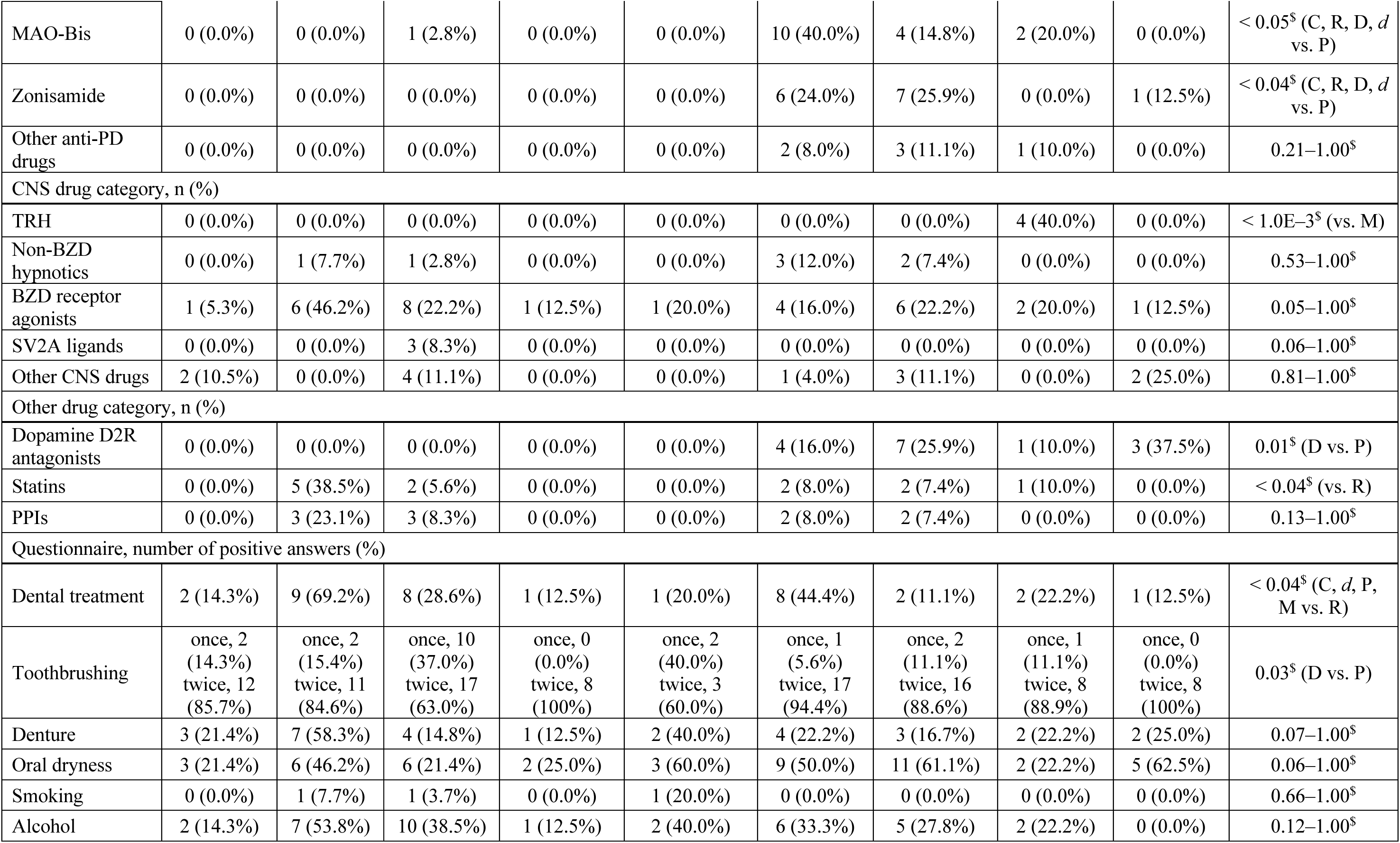
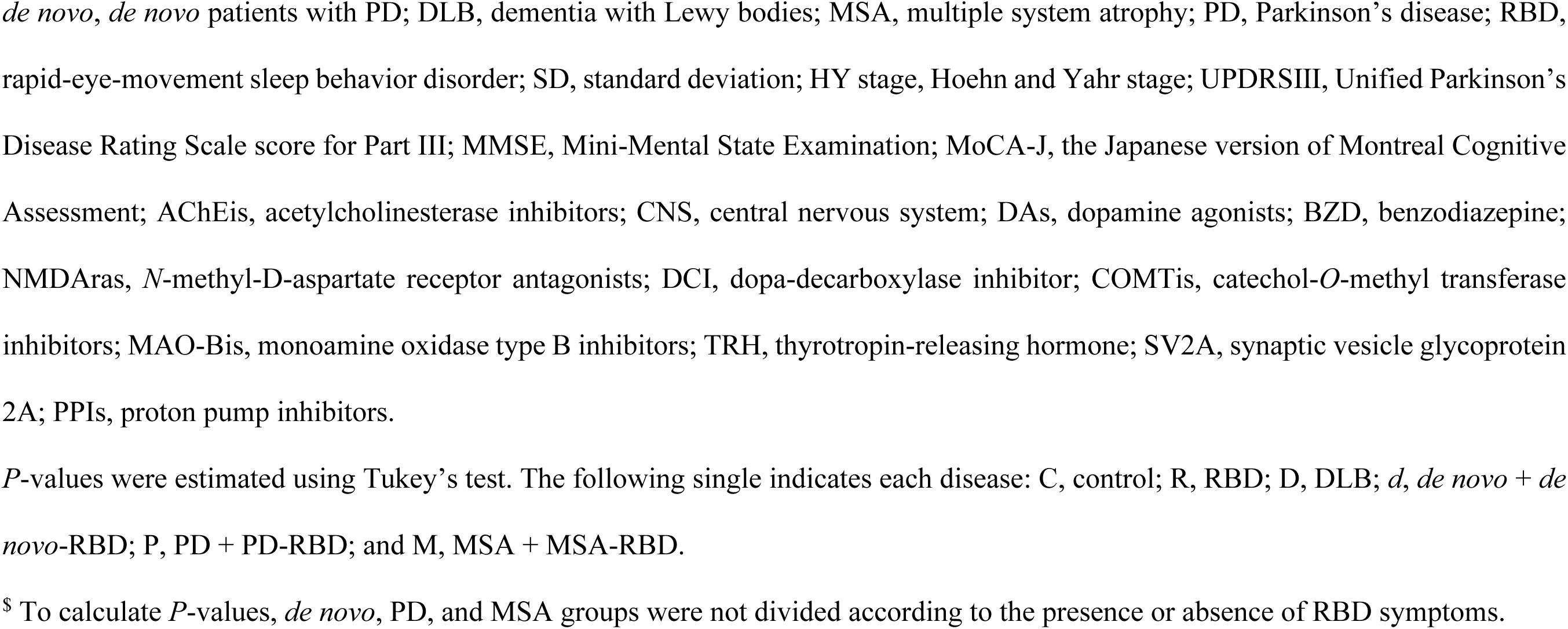
Demographics and the information on prescribed drugs in the primary cohort.

### Differential and shared microbial alteration and functional prediction across RBD and synucleinopathy groups

To explore microbiome alteration characteristics in the prodromal stage and synucleinopathies, we first defined age, sex, disease, anti-dementia and PD (amantadine HCl) drugs, statins, and proton pump inhibitors (PPIs) as covariates in MaAsLin2, according to the stepwise genus-level RDA (Supplementary Table 1) and our previous reports^21,27^. Multivariate analyses adjusted for these covariates revealed that the DLB and PD-RBD groups had significantly reduced Shannon diversity (evenness within an individual), whereas the *de novo* group showed increased diversity compared with both the control and RBD groups (*P*<0.05; Fig. 2a,b). We found that eight genera differed significantly between the RBD and control groups (*P*<0.05; Fig. 2a). The MSA group showed the most significant changes, with 10 and 8 genera compared with the control and RBD groups, respectively (*P*<0.05; Fig. 2a,b). *Atopobium* and *Alloprevotella* abundances increased significantly in seven and five disease groups, respectively, compared with the control group (*P*<0.05; Fig. 2a and Supplementary Table 2), whereas the abundance decreased, especially in the DLB and *de novo* groups, compared with the RBD group (Fig. 2b and Supplementary Table 3).

**Fig. 2.**
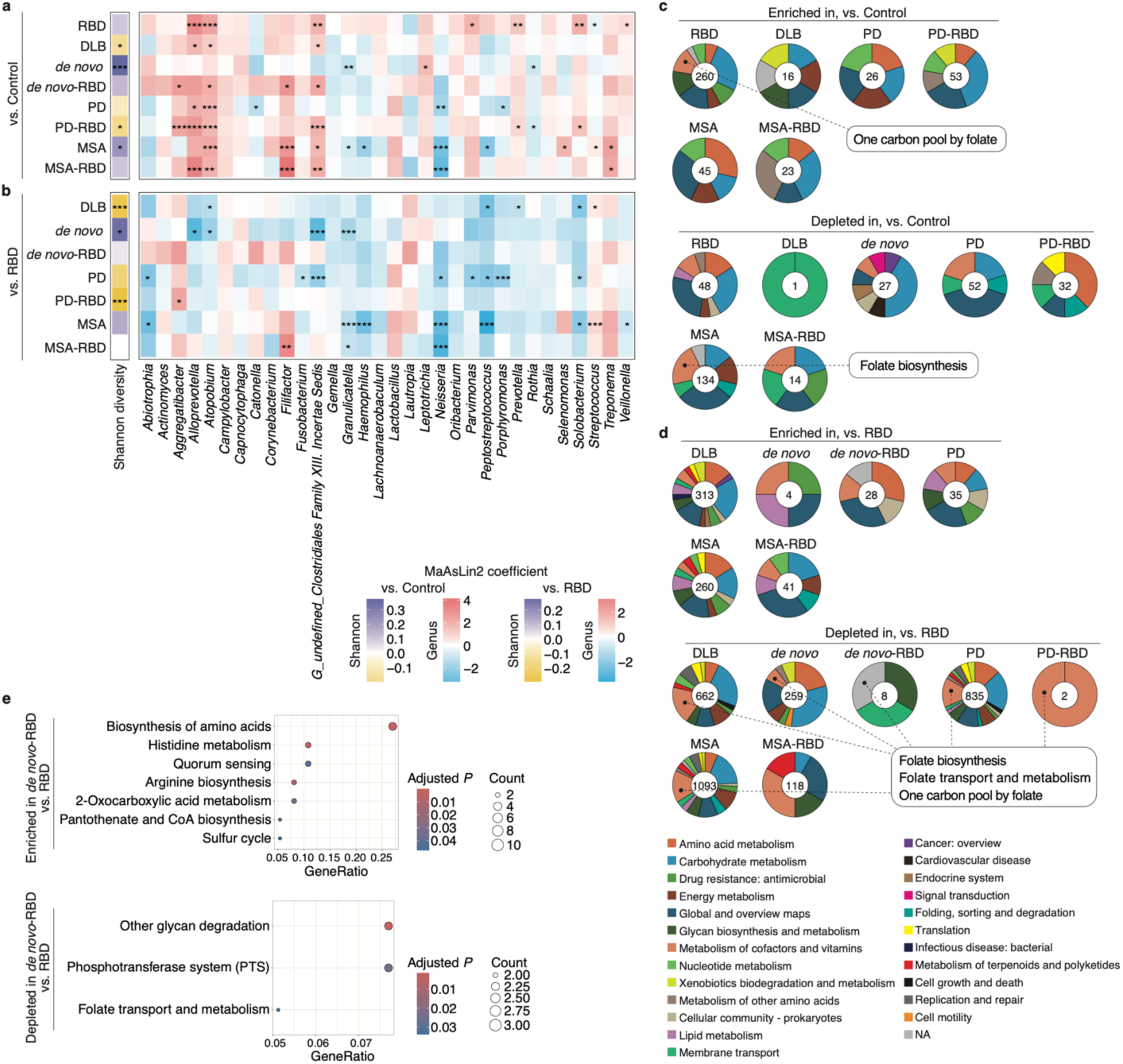
Differential and shared microbial alteration and functional prediction across the RBD and synucleinopathies. **a**,**b**, Heatmaps showing the enrichment and depletion of the top 32 most abundant genera (relative mean abundance >0.1%) and Shannon diversity scores between the control (**a**) or RBD (**b**) and each disease group (*n*=151). Statistical significance was determined through the MaAsLin2 package, incorporating age, sex, disease, AChEis, NMDAras (memantine and amantadine HCl), statins, and PPIs as random effects. \**P*<0.05; \*\**P*<0.01; \*\*\**P*<0.005. **c**,**d**, Pie charts showing the subcategory of PICRUSt2-predicted KEGG pathways that were significantly enriched and depleted between the control (**c**) or RBD (**d**) and each disease group (*n*=151). The numbers in the circles indicate the number of genes contained. The descriptions related to “folate” are shown in the figure. **e**, Bubble plots showing the description of significant KEGG pathways (**d**) with the highest GeneRatio (adjusted *P*<0.05) between the RBD and *de novo*-RBD groups. The adjusted *P*- values were corrected using the Benjamini–Hochberg method. Significant KEGG orthology terms were determined using the MaAsLin2 package, incorporating age, sex, disease, AChEis, NMDAras (memantine and amantadine HCl), statins, and PPIs as random effects (*P*<0.05). AChEi, acetylcholinesterase inhibitor; *de novo*, *de novo* patients with PD; DLB, dementia with Lewy bodies; KEGG, Kyoto Encyclopedia of Genes and Genomes; MSA, multiple system atrophy; NMDAra, *N*-methyl-D-aspartate receptor antagonist; PD, Parkinson’s disease; PPI, proton pump inhibitor; RBD, rapid-eye-movement sleep behavior disorder.

*Peptostreptococcus* and *Solobacterium* abundances decreased significantly in the DLB, PD, and MSA groups compared with the RBD group (*P*<0.05; Fig. 2b and Supplementary Table 3).

Next, we analyzed the predictive metabolic pathways associated with synucleinopathies using phylogenetic investigation of communities by reconstruction of unobserved states (PICRUSt2). The significantly enriched and depleted pathways in the disease groups compared with those in the control and RBD groups are shown in Fig. 2c,d and Supplementary Tables 4–7 (MaAsLin2, *P*<0.05). Numerous genes were included in the depleted pathways of the synucleinopathy groups compared with the RBD group, indicating large metabolic differences between the DLB, PD, MSA, and RBD groups (Fig. 2d). For example, “folate transport and metabolism” pathway was significantly depleted in the *de novo*-RBD group compared with the control group (adjusted *P*<0.05; Fig. 2e). Notably, folate-related pathways (folate biosynthesis, folate transport and metabolism, or one-carbon pool by folate) were commonly depleted in the synucleinopathy groups compared with the RBD group (Fig. 2c,d, and Supplementary Tables 4–7). These results imply that the salivary microbiome transiently changed (e.g., increased abundance) dramatically in patients with RBD compared with controls, subsequently showing patterns characteristic of individual synucleinopathies.

### Salivary microbiome profiles stratified RBD and synucleinopathies

We investigated the predictive microbial model for classifying prodromal and early PD stages based on salivary microbiome profiles. Based on the β-diversity analysis between subgroups of each disease (Fig. 1g), random forest classifiers were built in the primary cohort dataset without separating them according to RBD symptom presence or absence. We selected microbial features and generated predictive microbial models through 100 independent iterations using different random seeds, based on 20 repeats of 10-fold cross-validation (Supplementary Fig. 1a). The performance of the generated predictive model for the RBD and *de novo*+*de novo*-RBD groups was validated using an independent test cohort dataset, which was randomly partitioned into a training set (70%) and a test set (30%). The demographics of the test cohort are shown in Supplementary Table 8. The median area under the receiver operating characteristic curve (AUROC) and the area under the precision recall curve in the predictive models of the primary cohort for classifying each synucleinopathy from the control, RBD, or *de novo*+*de novo*-RBD groups are shown in Fig. 3a–f. We selected 56 and 60 species as microbial features based on their variable importance in the predictive model comparing RBD and *de novo*+*de novo*-RBD groups with all other groups, respectively (Supplementary Fig. 1b and 2). For the independent test cohort, the median AUROCs and 95% confidence intervals (CI) for classifying *de novo*+*de novo*-RBD from each group (control, DLB, PD+PD-RBD, and MSA+MSA-RBD) were 0.78 (0.76–0.81), 0.80 (0.79–0.84), 0.85 (0.84–0.89), and 0.86 (0.86–0.90), respectively (Fig. 3g). In the RBD predictive model versus that of each group (control, *de novo*+*de novo*-RBD, DLB, PD+PD-RBD, and MSA+MSA-RBD), the median values were 0.85 (0.82–0.87), 0.86 (0.84–0.89), 0.86 (0.84–0.89), 0.89 (0.88–0.91), and 0.94 (0.92–0.96), respectively (Fig. 3h). The AUROC in the RBD model versus control was higher than that in the *de novo*+*de novo*-RBD model versus control, which is consistent with the weighted UniFrac distance analysis results (Fig. 1h).

**Fig. 3.**
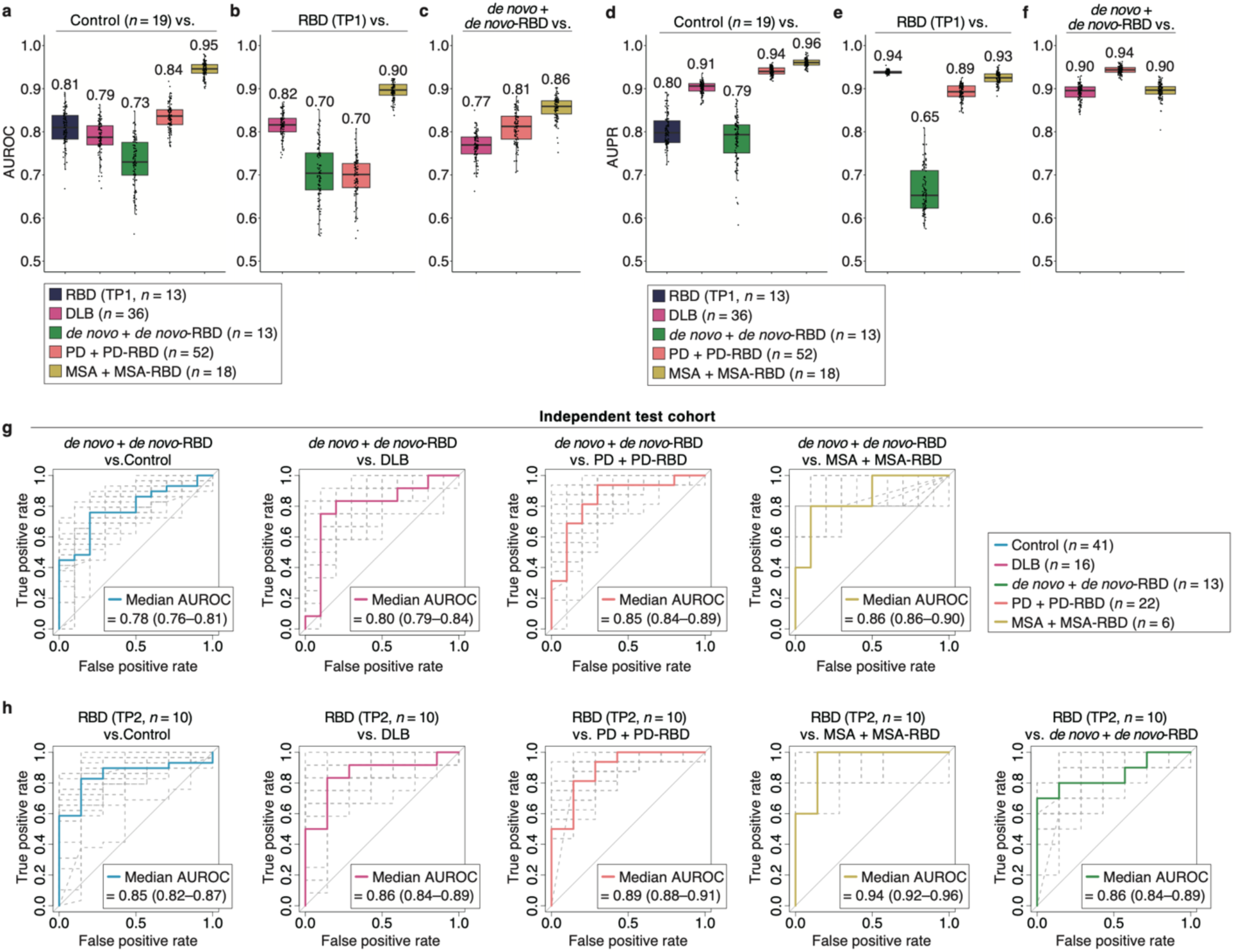
Predictive microbial models to stratify RBD or early PD stage from synucleinopathies. **a–f**, Graphs showing the area under the receiver operating characteristic curve (AUROC) and the precision recall curve (AUPR) in random forest classifiers generated by abundant 70 species (relative mean abundance >0.1%) of the primary cohort. The box plots present the median and 25th and 75th percentiles. The dots indicate the AUROC or AUPR of the single iterations. The number of participants is shown in the figure. **g**,**h**, Graphs showing the ROC curves of the independent test cohort for classifying the *de novo*+*de novo*-RBD (**g**) or RBD (**h**) from each disease group in the predictive microbial models constructed using microbial features selected by 100 iterations (Supplementary Figs. 1 and 2). The dashed gray lines represent the results from 25 resamplings, and each bold-colored line indicates the median. The number of participants is shown in the figure. *de novo*, *de novo* patients with PD; DLB, dementia with Lewy bodies; MSA, multiple system atrophy; PD, Parkinson’s disease; RBD, rapid-eye-movement sleep behavior disorder; TP, time point.

### Visualization of the salivary microbiome compositional manifold with pseudotime

Based on the results indicating the potential for RBD classification with high accuracy, we investigated whether the salivary microbiome serves as a prospective biomarker of progression from RBD to synucleinopathies. We applied pseudotime trajectory analyses to our salivary microbial data, as previously described^26^. First, we input the microbial abundance data at the species level for all samples (Fig. 1a), including the time-course samples collected longitudinally from the same individual, into a computational framework that enabled the identification of various low-dimensional roots (or branches) in the microbiome compositional space and the location of samples along these roots (Fig. 4a). Second, the framework labeled all samples with pseudotimes based on their distance from a predefined root sample. We set the control and RBD groups or the other groups as the root of preclinical or disease stages, respectively. Notably, in cases where the disease stage was the root, we applied a reversed pseudotime to converge on the root. Finally, we estimated disease progression based on the branches and analyzed the changes in bacterial abundance and functional predictions associated with the branches. We identified eight clusters and four branches in the microbiome compositional space that were related to information on the disease category and cohort (Fig. 4b). The characteristic pseudotimes were observed on the manifold, depending on the preclinical and disease stages (Fig. 4c,d). We showed all sample characteristics corresponding to the pseudotime of setting the control as the root and found that diverse pseudotime ranges and clusters were included in each disease category (Fig. 4e and Supplementary Table 9). We further identified four branches originating from cluster 6, containing most preclinical samples, based on between-cluster connectivity (Fig. 4f,g).

**Fig. 4.**
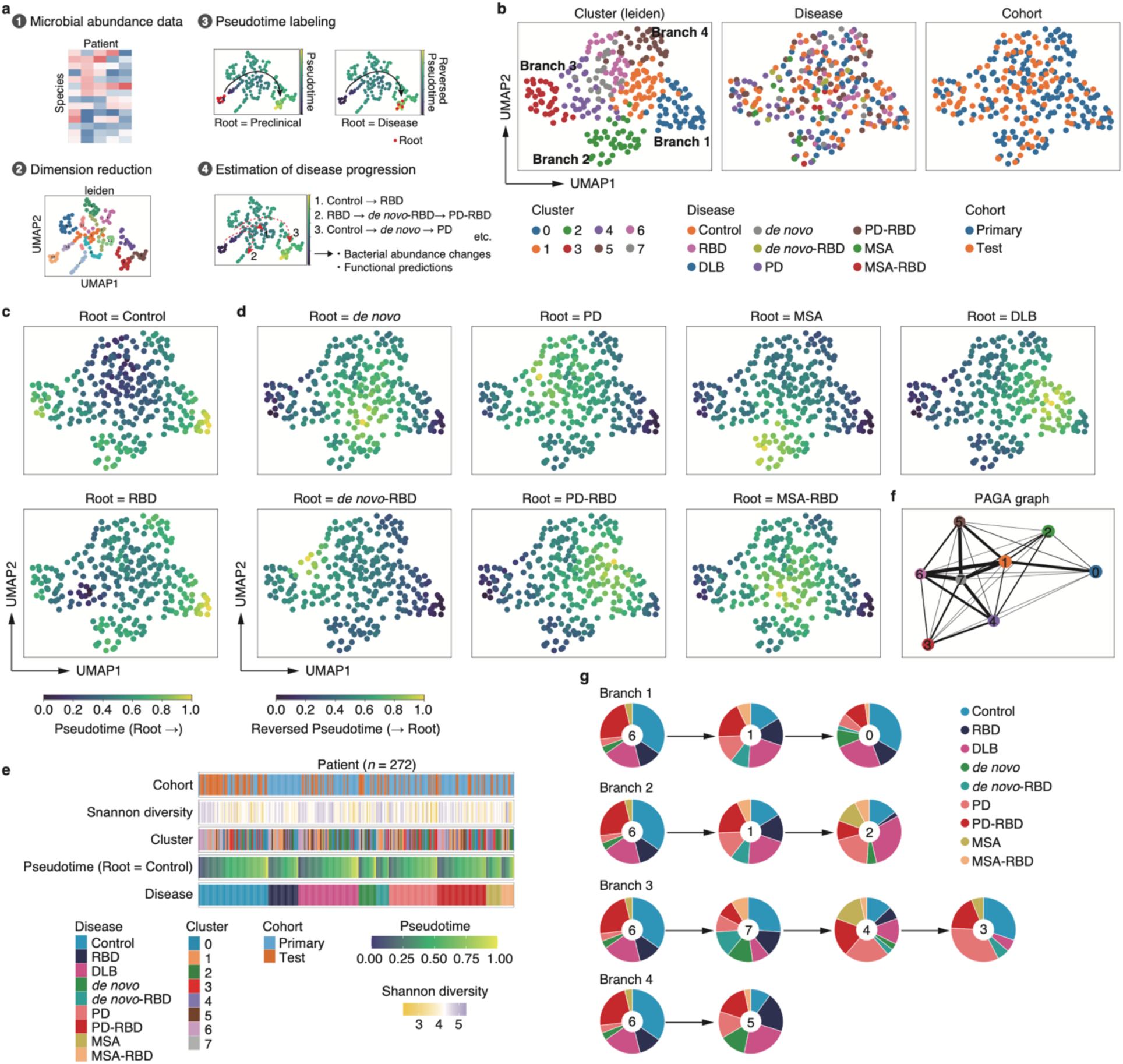
Pseudotime and branch of the salivary microbiome composition and its association with diseases. **a**, Framework of dimensionality reduction and pseudotime labeling using the partition-based graph abstraction (PAGA) algorithm. **b**, Graphs visualizing the microbiome compositional manifold (eight clusters and four branches) and its sample characteristics. **c**,**d**, Scatter plots showing the pseudotime (**c**) and reversed pseudotime (**d**) in the preclinical and disease stages as the root, respectively. **e**, Heatmaps showing the characteristics of each sample corresponding to the pseudotime of setting the control as the root. **f**, The PAGA graph represents the connectivity between partitions by associating nodes with each partition and quantifying the edges with weights. **g**, Pie charts showing each branch, including cluster and disease groups. *de novo*, *de novo* patients with PD; DLB, dementia with Lewy bodies; MSA, multiple system atrophy; PD, Parkinson’s disease; RBD, rapid-eye-movement sleep behavior disorder; UMAP, uniform manifold approximation and projection.

### Individual trajectories and pseudotimes associated with longitudinal analysis

To compare our pseudotimes and real-world longitudinal samples, we investigated the locations of individual longitudinal samples on the manifold. We presented plots that included longitudinal samples from 10 patients with iRBD, observing varying distances between samples for each individual (Supplementary Fig. 3a,b). No significant difference was observed in the change in pseudotime between the two sampling times (pseudotime Δt; Supplementary Fig. 3c). We also investigated the location of the longitudinal samples in 10 *de novo* patients, including *de novo*- RBD, which indicated changes in the sample position associated with anti-PD drug treatment (Supplementary Fig. 4a–c). We found that the pseudotime Δt differed significantly in the reversed pseudotimes of PD or PD-RBD as the root (*P*=0.048 and 0.029, respectively; Supplementary Fig. 4d,e). The increased pseudotime Δt observed when PD or PD-RBD was used as the root suggests that the post-treatment samples of *de novo* patients moved closer to the centroid of the PD or PD- RBD groups. These results imply that our pseudotime analysis aligns with real-world longitudinal samples.

### Shifted microbial abundances with pseudotime

To identify microbiome alterations associated with disease progression in RBD, we first examined the correlation between pseudotime and disease duration. Pseudotime (root=RBD) was positively correlated with disease duration in both RBD and *de novo*-RBD groups (Supplementary Fig. 5a). However, a negative correlation was observed in both PD-RBD and MSA-RBD groups. In our cohort, patients with iRBD who converted to PD or DLB within 1 year after sampling were defined as progressors, and the remaining patients with iRBD were defined as nonprogressors. Disease duration and pseudotime tended to increase in progressors compared to nonprogressors (Supplementary Fig. 5b,c), indicating that the pseudotime partially reflects disease progression within RBD. We observed changes in the normalized microbial abundance with pseudotime and identified 11 species that were significantly correlated (*P*<0.05; Supplementary Fig. 5d). For example, *Neisseria perflava* abundance exhibited a significant positive correlation, whereas that of *Streptococcus salivarius* exhibited a significant negative correlation (rho=0.58 and −0.59, respectively; *P*<0.005).

We further investigated the correlation between the reversed pseudotime and normalized microbial abundance in each disease group. The *de novo*, *de novo*-RBD, PD, PD-RBD, MSA, MSA-RBD, and DLB groups included significantly correlated 17, 25, 21, 19, 19, 12, and 20 species, respectively (*P*<0.05; Supplementary Fig. 6). Among the significantly correlated species, two, four, and seven species were shared between *de novo* and *de novo*-RBD, PD and PD-RBD, or MSA and MSA-RBD groups, respectively (Supplementary Fig. 6a−c).

### Distinct microbial alteration and functional prediction in branch 3 with a short pseudotime from prodromal to early PD stages

Finally, we investigated the features, including host factors, in each branch to clarify whether specific conditions contributed to manifold formation in the salivary microbiome composition, which is influenced by various factors. The demographics of the branches are presented in Supplementary Table 10. We found that the pseudotime (root=RBD) in the *de novo*, PD, or both groups in branch 3 was significantly shorter than that in other branches (*P*<0.031; Fig. 5a–c). Furthermore, the Shannon diversity was significantly lower in branch 3 than in other branches (*P*<0.019; Fig. 5d). To investigate the microbial features in branch 3, we compared the normalized genus-level abundance among the branches. We identified seven genera that significantly differed in branch 3 compared to the other branches (*P*<0.05; Fig. 5e). The abundances of *Streptococcus* and *Lactobacillus* were enriched in branch 3, whereas those of *Neisseria*, *Haemophilus*, *Peptostreptococcus*, and *Porphyromonas* were depleted, as shown by the normalized abundances on the manifold (Fig. 5f). Further multivariate analyses of the predictive metabolic pathways revealed that 17 and 32 pathways were significantly depleted and enriched in branch 3, respectively, compared to all other branches (adjusted *P*<0.05; Fig. 5g,h). Depleted pathways included nucleotide sugar biosynthesis, sulfur metabolism, and oxidative phosphorylation. Enriched pathways included amino acid biosynthesis, ribosome, nucleotide metabolism, and peptidoglycan biosynthesis.

**Fig. 5.**
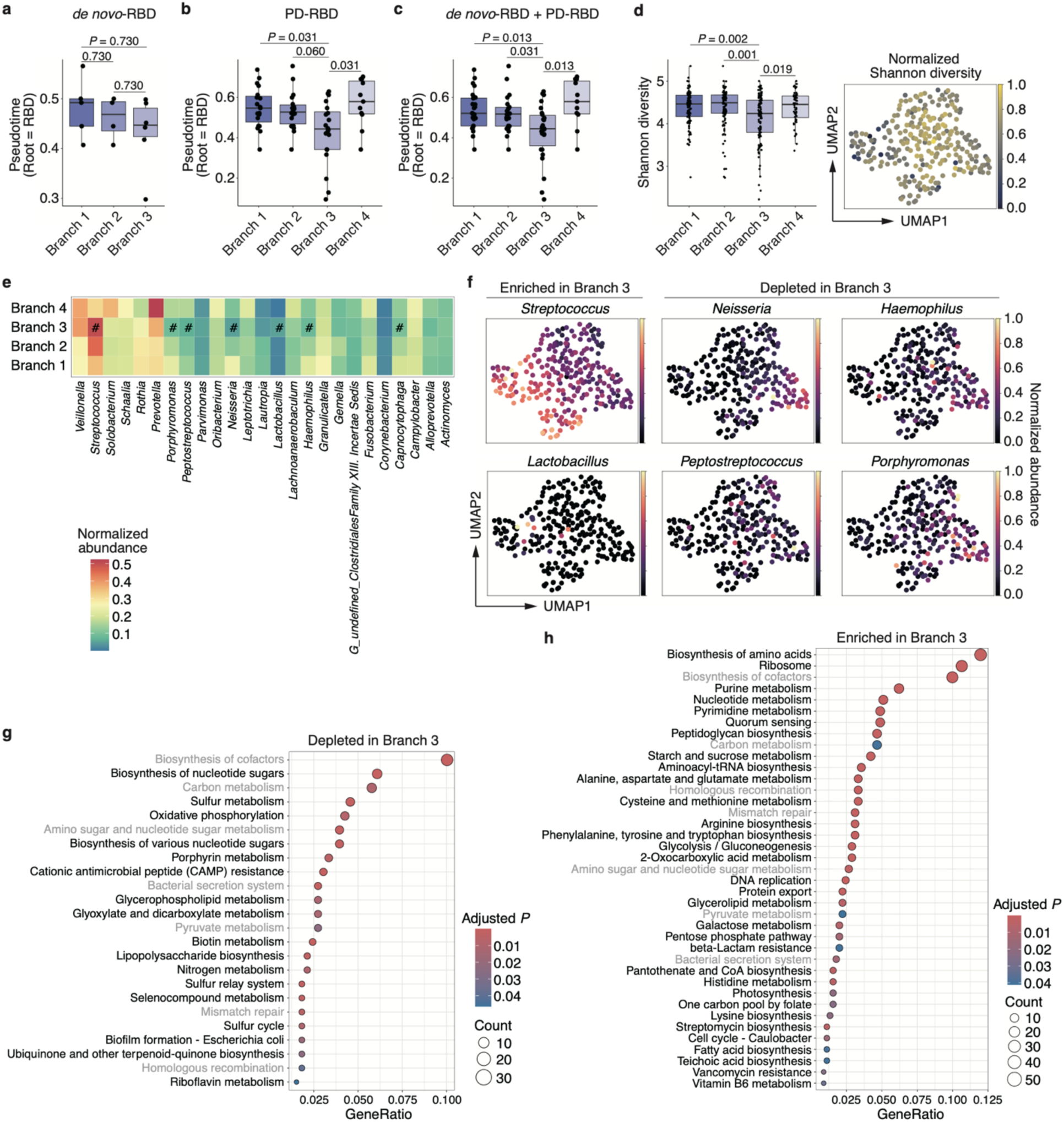
Distinct microbial alteration and functional prediction in branch 3 with a short pseudotime from prodromal to early PD stage. **a–c**, Graphs showing the pseudotime (RBD as the root) of *de novo*-RBD (**a**), PD-RBD (**b**), and *de novo*-RBD + PD-RBD (**c**) in each branch. The box plots present the median and 25th and 75th percentiles. The dots indicate each participant, and the distribution in each branch is as follows: *de novo*-RBD: branch 1=5, branch 2=4, and branch 3=6; PD-RBD: branch 1=19, branch 2=18, branch 3=20, and branch 4=11. Statistical significance was determined using the Wilcoxon rank-sum test with the Benjamini–Hochberg method (*P*<0.05). **d**, The left panel shows Shannon diversity by branch. Statistical significance was determined using the Wilcoxon rank-sum test with the Benjamini–Hochberg method (*P*<0.05). Right scatter plot where samples are colored according to the normalized Shannon diversity score. **e**, Heatmap showing the normalized abundance of the top 25 genera (relative mean abundance >0.1%) in each branch. The distribution of branches is as follows: branch 1=114, branch 2=110, branch 3=113, and branch 4=56. Statistical significance was determined using the Wilcoxon rank-sum test with the Benjamini–Hochberg method (*P*<0.05). ^#^The bacteria in branch 3 showed significant differences (*P*<0.05) compared to those in all other branches. **f**, Scatter plots where samples are colored according to the normalized abundance of representative genera significantly enriched and depleted in branch 3. **g**,**h**, Bubble plots showing the description of significant KEGG pathways with the highest GeneRatio (adjusted *P*<0.05) depleted (**g**) and enriched (**h**) in branch 3. The adjusted *P*-values were corrected using the Benjamini–Hochberg method. Significant KEGG orthology terms were determined using the Wilcoxon rank-sum test with the Benjamini–Hochberg method (*P*<0.05). The gray text indicates the shared KEGG pathways between enriched and depleted pathways. *de novo*, *de novo* patients with PD; KEGG, Kyoto Encyclopedia of Genes and Genomes; PD, Parkinson’s disease; RBD, rapid-eye-movement sleep behavior disorder.

To evaluate the host factors contributing to branch formation, we selected the variable importance using 25 independent iterations of the random forest classifier, based on 20 repeats of 10-fold cross-validation (Supplementary Fig. 7a). This classifier takes as input pseudotime (root=RBD), Shannon diversity score, age, sex, body mass index (BMI), and questionnaire responses (dental treatment, toothbrushing, denture, oral dryness, smoking, and alcohol consumption). Pseudotime, Shannon diversity, age, BMI, and oral dryness were identified as contributing factors for classifying branch 3 into at least two other branches (Supplementary Fig. 7b).

## Discussion

The main finding of this study was that RBD can be classified from non-neurodegenerative controls with high discriminative ability (AUC=0.85, 95% CI=0.82–0.87) and stratified from synucleinopathies, including early PD (AUC range: 0.86–0.94). Notably, the salivary microbiome composition of patients with iRBD was closer to that of synucleinopathies with RBD symptoms than that of synucleinopathies without RBD symptoms (Supplementary Fig. 8). We previously evaluated patients with iRBD using dopamine transporter single-photon emission computed tomography (DaT) and ^123^I-*meta*-iodobenzylguanidine (MIBG) myocardial scintigraphy, which assesses nigrostriatal dopaminergic function and cardiac sympathetic nerve degeneration, respectively^28^. The RBD cohort included 103 patients (95%) with decreased MIBG uptake; among them, 28% also showed decreased DaT uptake but 72% had normal DaT uptake. No patients with iRBD exhibited normal MIBG uptake with decreased DaT uptake. These results suggest that the RBD group in this study represents the body-first type^28^. Collectively, salivary dysbiosis in the prodromal stage is consistent with the idea that gut dysbiosis could predispose to the body-first type^2^. We further identified that a specific population with low diversity, enriched *Streptococcus*, and depleted *Neisseria* had a short distance from RBD to early PD by our pseudotime trajectory analyses (hereinafter “short pseudotime population”). This population was partially associated with host factors, such as BMI and oral dryness. These results suggest that RBD-to-PD progression may be accelerated in the short pseudotime population.

The results of the weighted UniFrac distance, Kyoto Encyclopedia of Genes and Genomes (KEGG) enrichment (the number of included genes), and pseudotime (the number of significant correlation species) analyses suggest that the effect of RBD symptoms on the salivary microbiome is strongly reflected in the *de novo* and PD groups, whereas in the MSA group, MSA symptoms might be pronounced independent of RBD presence. These results may be related to RBD being more likely to progress to PD or DLB than to MSA^2^. Whereas three metabolic pathways (pentose and glucuronate interconversions, carbon metabolism, and butanoate metabolism) were commonly enriched in patients with iRBD, PD-RBD, and MSA-RBD compared to controls, indicating the presence of metabolic pathways characteristic of patients with RBD symptoms. Butanoate metabolism produces butyrate, a short-chain fatty acid (SCFA), which mainly exerts positive effects, including acting as an energy source for epithelial cells and having anti-inflammatory and neuroprotective properties^29^. SCFA-producing gut bacteria depletion triggers intestinal hyperpermeability, immune activation, and enteric pathological α-synuclein aggregation in patients with PD^30–32^. Huang et al. found that SCFA-producing bacteria are already depleted in the gut microbiome of patients with RBD^15^; however, this depletion is controversial^12,33^. In contrast, high concentrations of SCFAs, especially butyrate, produced by oral bacteria, are associated with periodontitis^34,35^. These results suggest that bacterial functions depend on the sites or environments in which they exist. Although several studies have reported a relationship between salivary SCFAs and neurodegeneration^36,37^, the role of SCFA-producing oral bacteria in patients with synucleinopathies remains unknown.

Furthermore, specific metabolic pathways (folate biosynthesis, folate transport and metabolism, or one-carbon pool by folate) were commonly depleted in synucleinopathy compared to RBD. Serum depletion of folate, a vitamin, has been observed in patients with PD^38^. One proposed mechanism in animal experiments is that folate-related metabolic abnormalities trigger neuroinflammation, resulting in cell death^39,40^. Previous studies have reported a decrease in the vitamin biosynthesis in the gut microbiome of patients with PD^15,41,42^, consistent with our findings. However, vitamin deficiency in patients with PD is caused by reduced dietary intake or digestive disorders rather than gut microbiome metabolism changes^43^. Considering our previous results showing a low correlation between the gut and salivary microbiomes in patients with AD and PD^21^, the interaction between pathogenesis and the salivary microbiome in patients with RBD and synucleinopathies requires further investigation.

Salivary *Streptococcus* and *Lactobacillus* are more abundant in patients with PD than in controls^19^, which does not correspond with our multivariate analysis findings. This difference may be due to our multivariate analysis considering confounders such as anti-PD drugs that affect the salivary microbiome composition. However, our trajectory analyses revealed that the increase in the abundance of these two genera was observed in the short pseudotime population. These results could not be obtained by performing multivariate analyses between the disease categories.

*Streptococcus* is a dominant genus in the oral cavity, and *Lactobacillus* is associated with dental caries^44^. The overabundance of oral *Streptococcus* and pathogens, including *Lactobacillus*, is related to local or systemic inflammation and systemic diseases^45,46^. Interestingly, an increase in *Lactobacillus reuteri* abundance might augment α-synuclein secretion in the enteric nerves via neuron activation^19,47,48^. Regarding the depleted bacteria in the short pseudotime population, *Peptostreptococcus* species exert anti-inflammatory properties on the intestinal epithelial barrier^49^. The products (nitrite) of *Neisseria* and *Haemophilus,* nitrate-reducing bacteria, have beneficial effects in local and systemic diseases, such as reducing periodontitis-related pathogens and neuroinflammation or ameliorating cardiovascular disease by improving systemic blood circulation^50,51^. A beneficial interrelationship between *Neisseria* and *Haemophilus* has been suggested by the horizontal gene transfer that occurs between *Haemophilus* and *Neisseria meningitidis*^52^. Intriguingly, L’Heureux et al. proposed a potential intervention for cognitive impairment through the management of oral bacteria, including increasing N*eisseria* and *Haemophilus* abundance^51^. The nitrogen metabolism pathway was depleted, and the peptidoglycan biosynthesis pathway, related to inflammation, was enriched in the short pseudotime population. Our pseudotime trajectory analyses identified clusters or branches in the human salivary microbiome compositional space independent of individual disease categories, revealing host factors associated with these branches. These results were similar to those obtained using other algorithms on the gut microbiome dataset^25^, confirming the generalizability of our analysis. The proportion of participants who answered “yes” to oral dryness is as follows in branches 1, 2, 3, and 4: 33.3%, 38.3%, 52.9%, and 34.5%, respectively (Supplementary Table 10). Oral dryness perturbs oral/salivary microbiome resilience^53^, including the co-localization of non-oral bacteria, such as coliforms, and specific bacterium overgrowth, which might correspond to Shannon diversity reduction in the short pseudotime population. Our findings suggest that RBD-to-PD progression may be promoted in the short pseudotime population, which may be attributed to host factor-induced salivary microbiome alterations.

This study had some limitations. First, regarding the clinical sample, our cohort did not include naïve patients with DLB and iRBD at the time of onset. Almost all patients (86.1%) with DLB take anti-dementia drugs that affect the salivary microbiome. Patients seldom notice RBD symptoms; the symptoms are often identified by partners or others, making it challenging to determine the timing of onset and establish a large cohort including patients with iRBD at the early stage of onset. While early-onset iRBD analysis is indispensable for the development of early RBD diagnosis, our motivation was to stratify RBD from synucleinopathies and develop biomarkers for the RBD-to-PD progression. Considering that RBD-to-PD progression often takes 10 years, our cohort was well-matched in terms of motivation. In the prediction model, the *de novo*, PD, and MSA groups were not divided into two subgroups based on RBD symptom presence to avoid a small sample size. Moreover, almost all participants were diagnosed based on clinical and radiological features without neuropathological confirmation; it is unknown whether α-synuclein aggregates are present in the salivary gland or oral mucosa. Thus, we could not rule out patients with false clinical diagnoses. These clinical limitations may have affected the random forest classifier performance. Regarding technical limitations, previous trajectory analyses of the microbiome have employed large-scale datasets consisting of thousands of individuals based on publicly available data^25,26^. However, to our knowledge, no previous comprehensive study on the salivary microbiome has analyzed and publicly shared data on RBD and synucleinopathies considering RBD symptoms. Additionally, the current understanding of host factors remains incomplete. We recognize that our metadata collected from questionnaires may not fully reflect individual oral health, including periodontitis. Further investigations using comprehensive trajectory analysis, incorporating precise oral health information and other confounders (e.g., socioeconomic status or diet), are required to better understand the relationship between the salivary microbiome and synucleinopathies. Lastly, this study is limited to microbiome compositional data and functional predictions from 16S rRNA gene amplicon sequencing, which restricts our ability to identify species and strains accurately; metagenomics or long-read sequencing is needed in future studies to decipher salivary microbiome function in synucleinopathy pathogenesis.

Although the salivary microbiome-body-brain interaction mechanism in synucleinopathies remains elusive, harnessing the salivary microbiome holds the potential to stratify synucleinopathies, including RBD, which could lead to more effective interventions and the development of disease-modifying strategies.

## Methods

### Study design

We enrolled participants from two Japanese cohorts. The primary cohort consisted of patients with iRBD, DLB, PD, and MSA recruited from two hospitals located in Tokyo, a metropolitan area: Juntendo University Hospital (Tokyo, Japan) and Memory Clinic Ochanomizu (Tokyo, Japan). 62 patients with PD have been included in our previous study^21^. Patients with iRBD were recruited from our previous cohort^28^: Japan Parkinson’s Progression Markers Initiative (J-PPMI). The J-PPMI study, a prospective cohort, was conducted at multiple centers (National Center of Neurology and Psychiatry, Juntendo University, Kyoto University, Osaka University, and Nagoya University). The independent test cohort consisted of patients with DLB, PD, and MSA recruited from two hospitals located in a suburban area outside Tokyo: Juntendo University Urayasu Hospital (Chiba, Japan) and Memory Clinic of Toride (Ibaraki, Japan). The diagnosis was based on standard criteria (Movement Disorder Society [MDS]-sponsored PD clinical criteria^54^, MDS- sponsored MSA clinical criteria^55^, or definitions and guidelines provided by the DLB Consortium^56^), as previously described^10^. Patients with iRBD were examined using polysomnography as previously described^28^. In the *de novo*-RBD, PD-RBD, and MSA-RBD subgroups, RBD symptoms preceded motor symptoms. The control group consisted of individuals who exhibited no cognitive impairment based on cognitive function tests (Mini-Mental State Examination [MMSE] or the Japanese version of the Montreal Cognitive Assessment [MoCA-J]) and those without any neurodegenerative diseases in each cohort. Six participants taking anti-dementia drugs, despite having no cognitive impairment, were excluded from the control group (Fig. 1a). The overall exclusion criteria included oral diseases such as oral cancer and oral cenesthopathy (Fig. 1a). In patients with DLB, we excluded individuals with parkinsonism. We could not obtain precise clinical information on disease duration and RBD presence in patients with DLB. The minimum number of necessary samples for each experimental group was estimated based on our previous studies^10,21,27^.

For longitudinal analyses, we repeatedly collected saliva samples every 6 months from the initial sampling (Fig. 1a). Ten and three patients with iRBD were sampled twice and three times, respectively. Ten *de novo* patients with PD were sampled twice. Notably, four patients with iRBD progressed to synucleinopathies within 1 year of the last sampling.

The following metadata were collected: age, sex, BMI, clinical scores (MMSE, MoCA-J, Hoehn and Yahr [H&Y] scale, and MDS Unified PD Rating Scale score for Part III), prescription drugs, and questionnaire responses, as previously described^21,27^. A detailed questionnaire was used to assess smoking frequency (never or smoking) and alcohol consumption (regular [three or more times per week] or not regular). Dental treatment was divided into two scales: currently undergoing treatment and not undergoing treatment. Toothbrushing frequency was scored as once or more than twice. Participants were categorized into two groups in terms of denture usage: users and non-users. Oral dryness was self-reported as the presence or absence of a daily sensation of dryness. This study was approved by the Ethics Committee of Juntendo University School of Medicine (approval numbers H19-0244 and U20-0025). All participants or their legal guardians provided written informed consent to participate in this study. All procedures were performed in accordance with the principles of the Declaration of Helsinki.

### Classification of prescribed drugs

Drugs were classified based on their modes of action, as previously described^21,27^. Anti-dementia drugs included acetylcholinesterase inhibitors (AChEis) and *N*-methyl-D-aspartate receptor antagonists (NMDAras). We categorized anti-PD drugs into eight groups based on their mode of action: NMDAra (amantadine HCl), anticholinergics, levodopa/dopa-decarboxylase inhibitors, non-ergot-dopamine agonists (DAs), catechol-*O*-methyl transferase inhibitors, monoamine oxidase type B inhibitors, zonisamide, or others (including ergot-DAs, adenosine A2A receptor antagonists, and droxidopa). CNS drugs other than anti-dementia and PD drugs comprised thyrotropin-releasing hormone, non-benzodiazepine (BZD) hypnotics, BZD receptor agonists, synaptic vesicle glycoprotein 2A ligands, and other CNS drugs (including orexin receptor antagonists, selective serotonin reuptake inhibitors, noradrenergic and specific serotonergic antidepressants, tricyclic/tetracyclic antidepressants, melatonin receptor agonists, multi-acting receptor-targeted antipsychotics, aripiprazole, and lamotrigine). From our previous studies^21,27^, we selected three drug categories that may affect salivary microbiome composition: dopamine D2 receptor antagonists, statins, and PPIs. The proportions of these drugs are listed in Table 1 and Supplementary Table 8.

### Sample collection and 16S rRNA gene amplicon sequencing

We collected saliva samples during the daytime (9 a.m.–4 p.m.) using a sterilized sputum container (DE2000; Eiken Chemical Co., Tokyo, Japan); the samples were kept at 4℃, frozen in liquid nitrogen within 8 h after sampling, and stored at −80℃ until DNA extraction. We excluded participants who took antibiotics within 1 month before sampling, as previously described^57^. Sampling, freezing, and DNA extraction from frozen samples were performed using the enzymatic lysis method as previously described^57^. Briefly, we amplified the V1-V2 region of the 16S rRNA gene by polymerase chain reaction using the following primer sets: forward 27Fmod (5′- AATGATACGGCGACCACCGAGATCTACACxxxxxxxxACACTCTTTCCCTACACGACGC TCTTCCGATCTagrgtttgatymtggctcag-3′) and reverse 338R (5′- CAAGCAGAAGACGGCATACGAGATxxxxxxxxGTGACTGGAGTTCAGACGTGTGCTCT TCC GATCTtgctgcctcccgtaggagt-3′). Primer sequences included the Illumina Nextera adapter sequence and a unique 8-bp index sequence (indicated by xxxxxxxx) for each sample. Thermal cycling was performed on a 9700 PCR system (Life Technologies, Carlsbad, CA, USA) using Ex Taq polymerase (Takara Bio, Tokyo, Japan) with the following cycling conditions: initial denaturation at 96°C for 2 min; 25 cycles of denaturation at 96°C for 30 s, annealing at 55°C for 45 s, and extension at 72°C for 1 min; and final extension at 72°C. The amplicons were purified using AMPure XP magnetic purification beads (Beckman Coulter, Brea, CA, USA) and quantified using the Quant-iT PicoGreen dsDNA Assay Kit (Life Technologies, Japan). The amplicon pools were sequenced using the Illumina MisSeq Platform (2×300 bp), according to the manufacturer’s instructions.

### Data processing and microbiome compositional analysis

We used the analysis pipeline for MiSeq-barcoded amplicon sequencing, as described in previous studies^21,27^. Primer sequences were trimmed from the paired-end 16S rRNA gene amplicon using Cutadapt v.4.1–1. Amplicon sequence variants (ASVs) were constructed from the trimmed reads by removing primers and possible chimeric reads using the DADA2 R package v.1.18.0. We used 10,000 filter-passed reads per sample of high-quality ASVs and deposited them in the DDBJ/GenBank/EMBL database. The taxonomic assignment of ASVs was determined by similarity searches against the National Center for Biotechnology Information RefSeq database downloaded on January 8, 2020, using the GLSEARCH program.

Shannon diversity was calculated using the vegan R package v2.6−4 with the statistical programming language R, version 4.0.3 (2020-10-10). To determine the dissimilarity (distance) between each pair of samples, weighted UniFrac (considering both the presence/absence and relative abundance of species) distance analyses were performed as described in previous studies^21^. The β-diversity was obtained via permutational multivariate analysis of variance using weighted UniFrac distance, and *P*-values were adjusted using the Benjamini–Hochberg method.

### Multivariate analysis

Multivariate analyses were performed as previously described^21,27^. To estimate the linear cumulative and individual effect sizes of confounding variables that contribute to microbiome composition, we performed stepwise RDA based on the weighted UniFrac distance using the ordiR2step function (default direction=both) of the vegan R package v2.6−4. Individual adjusted *R*^2^ refers to the explained variance when the most influential variable is maximized in the first step involving all microbiome covariates. Differential microbial abundance analysis was performed via the microbiome multivariable associations with the linear model (MaAsLin2^58^) R package v1.10.0, using generated linear and mixed models (default model: min abundance=0.00; min prevalence=0.10; max significance=0.25; normalization=TSS; transformation=LOG). Based on the results of the stepwise RDA and our previous studies^21,27^, we selected age, sex, AChEis, NMDAras (memantine and amantadine HCl), statins, PPIs, and disease categories as random effects.

### Functional prediction

We used the PICRUSt2 software (v2.6.0.99) to predict the function of the microbiome community, as previously described^59^. Briefly, raw ASV count data were imported and processed through the PICRUSt2 pipeline using default parameters, and the aligned ASVs were placed into a reference tree to infer the gene family copy numbers. We obtained the abundance of KEGG orthology terms using the PICRUSt2 pipeline. Significant KEGG orthology terms were determined using the MaAsLin2 R package v1.10.0 (*P*<0.05), incorporating age, sex, AChEis, NMDAras (memantine and amantadine HCl), statins, PPIs, or disease categories as random effects. To identify significantly enriched and depleted predictive metabolic pathways (adjusted *P*<0.05), KEGG enrichment analysis was conducted using the clusterProfiler R package v4.4.4.

### Random forest classification and microbial feature selection

We performed random forest classification using the randomforest R package v4.7−1.1 and selected the microbial features, as described in previous studies^15,60^. The framework of random forest classification is shown in Supplementary Fig. 1. In the prediction model, *de novo*, PD, and MSA groups were not divided based on the presence of RBD symptoms, to avoid sample size reduction. First, to generate predictive microbial models using the primary cohort dataset, we performed 20 rounds of 10-fold cross-validation (inner loop) to discriminate the control group from the RBD, DLB, *de novo*, PD, and MSA groups. Likewise, cross-validations were performed to discriminate the RBD or *de novo* groups from the DLB, *de novo*, PD, and MSA groups or the DLB, PD, and MSA groups. Our motivation was to develop an early diagnostic method for RBD and early PD; therefore, we generated predictive microbial models for the RBD and *de novo* groups. To obtain microbial features (mean decrease in Gini) in the final predictive microbial models, 100 independent iterations of each inner loop were performed with different random seeds. Finally, the performance (median AUC and 95% CIs) of the microbial models for predicting the RBD and *de novo* groups was examined using 25 resamplings from the independent test cohort dataset, which was randomly partitioned into a training (70%) and a test (30%) set.

### Computational framework for detecting the manifold in the microbiome compositional space

We employed the framework provided by Tsamir-Rimon et al. to detect the manifold in the compositional space of the human microbiome^26^. The framework, consisting of the partition-based graph abstraction algorithm^24^, a general approach in the field of cell biology, includes default methods for dimensionality reduction, such as PCA and uniform manifold approximation and projection. First, the framework received microbial abundance data at the species level (relative mean abundance >0.05%) for all samples, including those in the longitudinal study. We adjusted the algorithm parameters (n_neighbors and n_psc) to form a distinct manifold following the pipeline provided by Tsamir-Rimon et al (https://github.com/borenstein-lab/vaginal_microbiome_manifold). All the samples were then ordered along the manifold and assigned pseudotimes using an extension of the diffusion maps. The diffusion map requires a predefined group, denoted as root samples, to represent the beginning of the manifold. Pseudotime is calculated using a random-walk-based diffusion pseudotime algorithm, resulting in a quantitative measure of progress through the biological process. Notably, in the present study, the predefined group denoted as root samples was divided into preclinical (control and RBD) and disease (*de novo*, *de novo*-RBD, PD, PD-RBD, MSA, MSA-RBD, and DLB) stages. Generally, healthy individuals have diverse microbial communities, which may become skewed in a specific direction (dysbiosis) depending on the disease. Thus, in cases where the disease stage was the root, we applied a reversed pseudotime to converge on the root (diseases). Finally, we estimated the branches starting from cluster 6, which contained the most preclinical samples based on clinical disease progression and connectivity between partitions (or clusters), by associating nodes with each partition and quantifying the edges with weights.

### Differential abundance analysis on the manifold

The four branches had overlapping clusters. The demographics of each branch that allowed sample overlapping are shown in Supplementary Table 10. The microbial abundance and Shannon diversity scores were normalized for manifold mapping. A significant difference in microbial abundance was obtained using the Wilcoxon rank-sum test with the Benjamini–Hochberg method. Spearman’s correlation coefficient analysis revealed a significant correlation between normalized microbial abundance and pseudotime or reversed pseudotime. For functional prediction, the KEGG orthology terms that changed significantly in branch 3, compared with all other branches, were determined using the Wilcoxon rank-sum test with the Benjamini–Hochberg method. To identify significantly enriched and depleted predictive metabolic pathways in branch 3 (adjusted *P*<0.05), we performed KEGG enrichment analysis using the clusterProfiler R package v4.4.4.

### Estimation of host factors contributing to branch formation

We performed random forest classification, incorporating potential host factors, using the randomforest R package v4.7−1.1. Variable importance (mean decrease in Gini) was selected from pseudotime (root=RBD), Shannon diversity score, age, sex, BMI, and questionnaire responses (dental treatment, toothbrushing, denture, oral dryness, smoking, and alcohol consumption) by 25 independent iterations of cross-validation (20 repeats of a 10-fold method).

### Statistical analyses

All statistical analyses were performed using the statistical programming language R version 4.3.3 (2024-02-29). Statistical significance was determined using Tukey’s test, the Wilcoxon rank-sum test via the Benjamini–Hochberg method, or the default methods in the following packages or analyses: stepwise RDA, MaAsLin2, and KEGG enrichment analysis. *P*-values or *Q*-values (or *FDR*) below 0.05 were considered significant.

## Data availability

The bacterial 16S rRNA sequencing data generated for this study were deposited in the DDBJ/GenBank/EMBL databases (accession numbers: DRA016579 and DRX705288- DRX705497 [available upon publication]). Clinical data, except those within this article, are not available in a public repository or supplementary material to protect the privacy and confidentiality of the study participants. Requests for clinical data can be directed to the corresponding authors and will be reviewed by the Ethics Committee of the Juntendo University School of Medicine. All shared data were de-identified.

## Code availability

Source codes used in this study are available from GitHub (available upon publication).

## Acknowledgements

We acknowledge all the participants. We are grateful to the medical staff in the Department of Neurology at Juntendo University Hospital, Juntendo University Urayasu Hospital, and the Memory Clinic. We thank Misaki Ouchida for creating the schematic illustrations. This work was supported by the Japan Society for the Promotion of Science (JSPS)/The Ministry of Education, Culture, Sports, Science, and Technology (MEXT), KAKENHI Grant-in-Aid for Early-Career Scientists (grant numbers 21K15888 and 24K17850 to D.H.), and the Japan Agency for Medical Research and Development (AMED) (21wm0425015 to T.H. and 21dm0207070 to N.H.). We also acknowledge financial support from the ITOCHU Chemical Frontier Corporation, Japan; Otsuka Holdings Co., Ltd., Japan; Meiji Seika Pharma Co., Ltd., Japan; and the Research and Development Board at Juntendo University.

## Author contributions

D.H., C.A., and N.H. designed and conceptualized the study. D.H. performed the overall data analysis. H.M., R.K., and W.S. conducted the 16S rRNA gene amplicon sequencing and data processing using an analysis pipeline. H.T.A., T.O., T.H., K.Y., and T.A. performed clinical examinations and provided clinical information. D.H., Y.M., and Y.N. recruited participants and completed the questionnaires. W.S., C.A., and N.H. supervised the study. D.H. wrote the manuscript with input from all the authors. All the authors have read and approved the final draft of this manuscript.

