## Supplemental information for "Leveraging the salivary microbiome profile to stratify REM sleep behavior disorder and synucleinopathies"

**Supplementary information**

**Supplementary Tables**

Supplementary Table 1. Explained variance in stepwise redundancy analysis at the genus level.

Supplementary Table 2. MaAsLin2 coefficients, adjusted for confounders (age, sex, AChEis, NMDAras [memantine and amantadine HCl], statins, and PPIs), in the RBD and synucleinopathies groups compared with the control group.

Supplementary Table 3. MaAsLin2 coefficients, adjusted for confounders (age, sex, AChEis, NMDAras [memantine and amantadine HCl], statins, and PPIs), in the synucleinopathies groups compared with the RBD group.

Supplementary Table 4. Significantly enriched KEGG orthology terms in the RBD and synucleinopathies groups compared with the control group.

Supplementary Table 5. Significantly depleted KEGG orthology terms in the RBD and synucleinopathies groups compared with the control group.

Supplementary Table 6. Significantly enriched KEGG orthology terms in the synucleinopathies groups compared with the RBD group.

Supplementary Table 7. Significantly depleted KEGG orthology terms in the synucleinopathies groups compared with the RBD group.

Supplementary Table 8. Demographics and the information on prescribed drugs in the test cohort.

Supplementary Table 9. Sample characteristics corresponding to the pseudotime of setting the control as the root.

Supplementary Table 10. Demographics and questionnaire responses in each branch.

### Supplementary figures

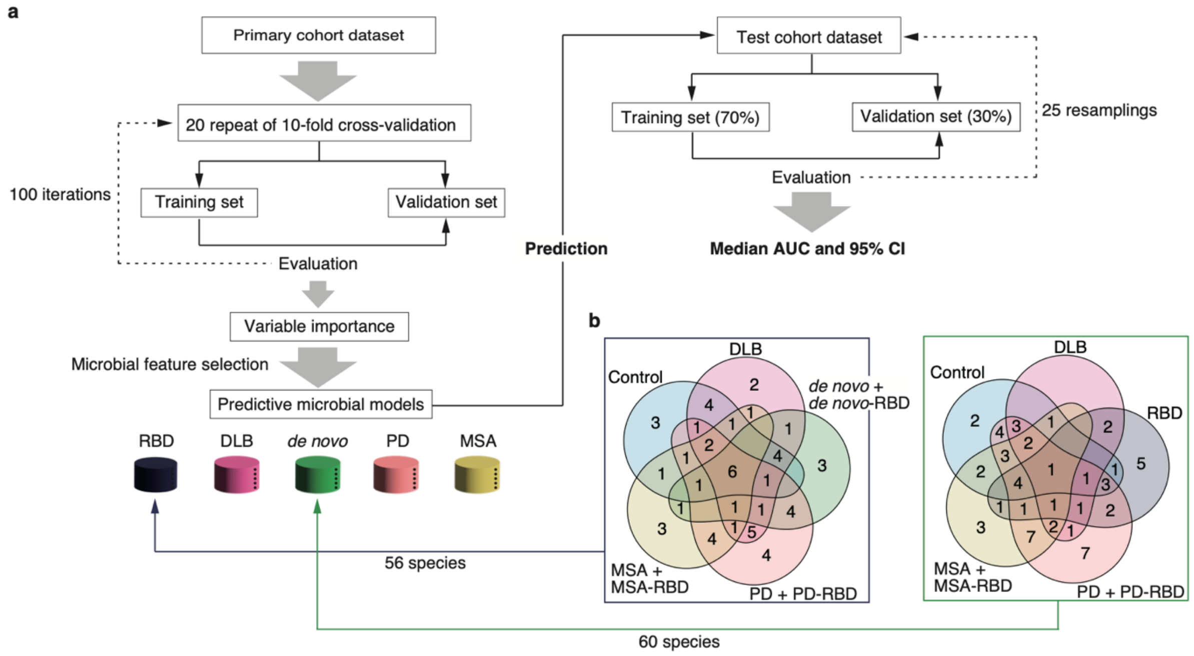


**Supplementary Fig. 1. Framework for constructing random forest models.**

**a**, For each random forest classification model (RBD, DLB, *de novo*+*de novo*-RBD, PD+PD-RBD, or MSA+MSA-RBD vs. control; DLB, *de novo*+*de novo*-RBD, PD+PD-RBD, or MSA+MSA-RBD vs. RBD; and DLB, PD+PD-RBD, or MSA+MSA-RBD vs. *de novo*+*de novo*-RBD), the primary cohort dataset (*n*=151) was randomly partitioned based on 20 rounds of 10-fold cross-validation (inner loop). The final predictive microbial models, incorporating the selected microbial features (mean decrease in Gini), were generated using 100 independent iterations of each inner loop with different random seeds. The performance (median AUC and 95% CIs) of the microbial models for predicting the prodromal (RBD) and early PD (*de novo*+*de novo*-RBD) stages was examined in the independent test cohort dataset (*n*=98), partitioned randomly into a training (70%) and a test (30%) set. **b**, The number of selected microbial features (Supplementary Fig. 2) in each random forest model for classifying prodromal and early PD stages.

AUC, area under the curve; CI, confidence interval; *de novo*, *de novo* patients with PD; DLB, dementia with Lewy bodies; MSA, multiple system atrophy; PD, Parkinson's disease; RBD, rapid-eye-movement sleep behavior disorder.

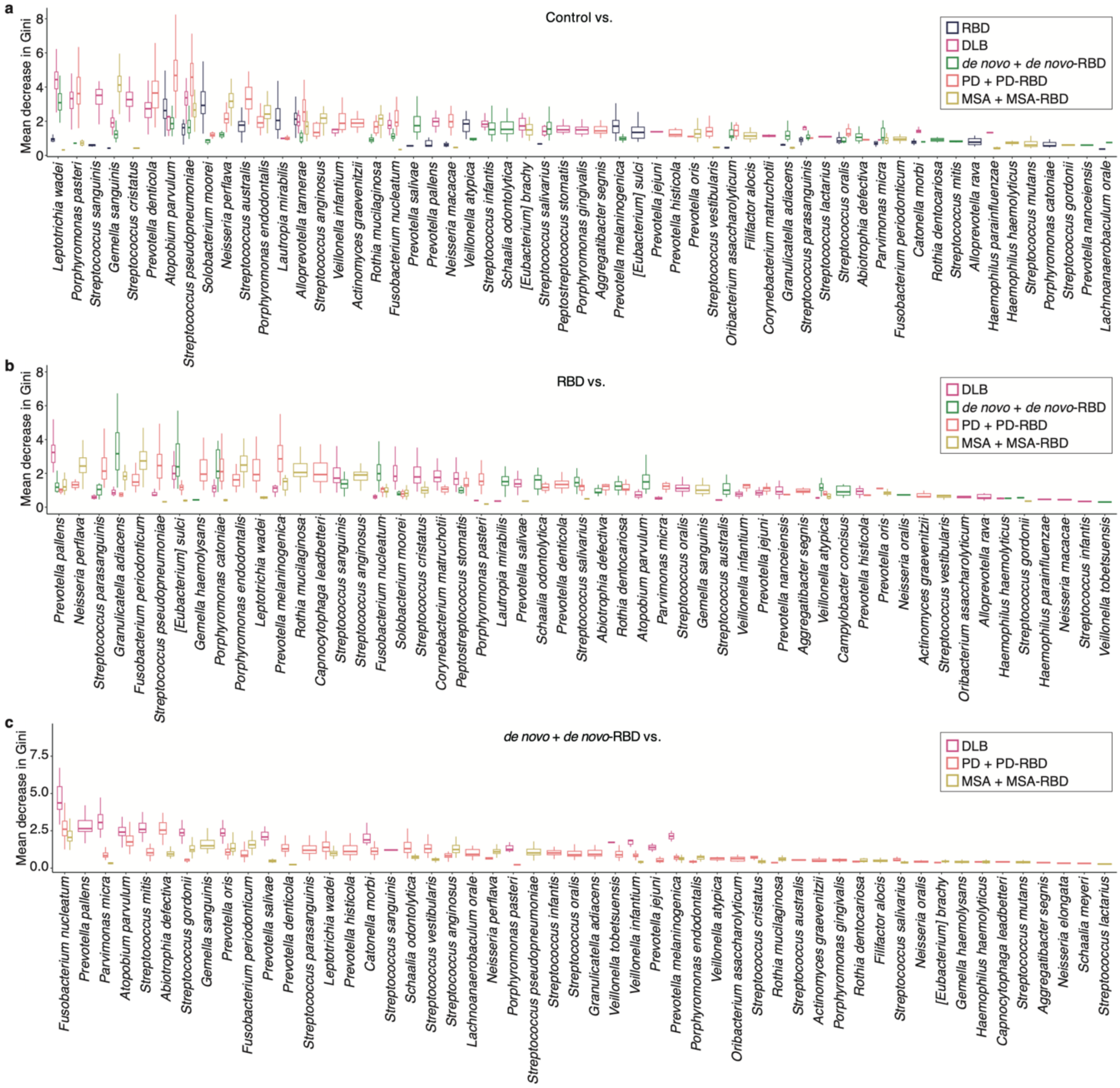


**Supplementary Fig. 2. Variable importance of random forest classifiers.**

**a–c**, The mean decrease in the Gini index, representing variable importance, was obtained from 100 independent iterations of an inner loop of the primary cohort dataset. The random forest classifiers take as input microbial data (70 species with a relative mean abundance of >0.1%). *de novo*, *de novo* patients with PD; DLB, dementia with Lewy bodies; MSA, multiple system atrophy; PD, Parkinson’s disease; RBD, rapid-eye-movement sleep behavior disorder.

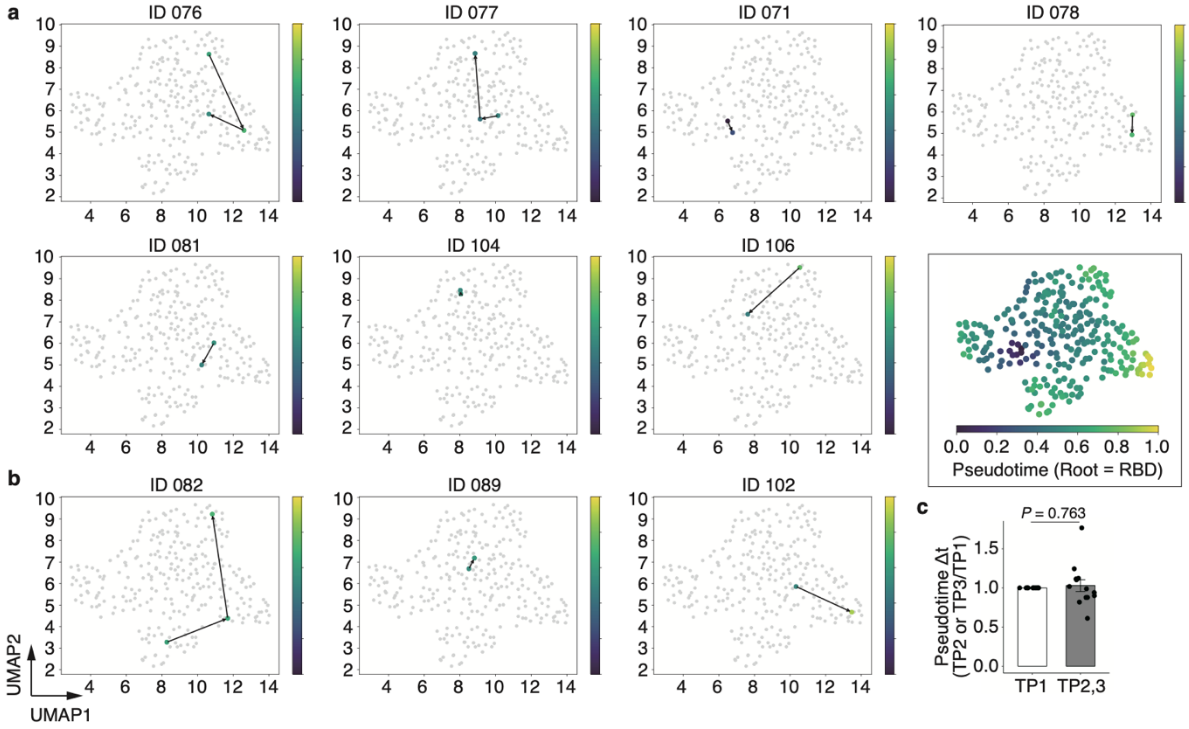


**Supplementary Fig. 3. Trajectory analysis with pseudotime in representative patients with RBD.**

**a**,**b**, Scatter plots showing the trajectory of the longitudinal analysis (twice or three times) in patients with RBD on the manifold with pseudotime. The gray dots represented the visualization of the microbiome compositional manifold, and colored dots indicated the locations of each sample from patients who have developed synucleinopathies (**b**) or still have RBD (**a**). Black arrows indicate the direction of the sampling time. **c**, Graph showing changes (Δt) in pseudotime (RBD as the root) between two sampling times (TP1, $n$=10; and TP2,3, $n$=13). Dots indicate each individual. Statistical significance was determined using the Wilcoxon rank-sum test ($P$<0.05).

RBD, rapid-eye-movement sleep behavior disorder; TP, time point.

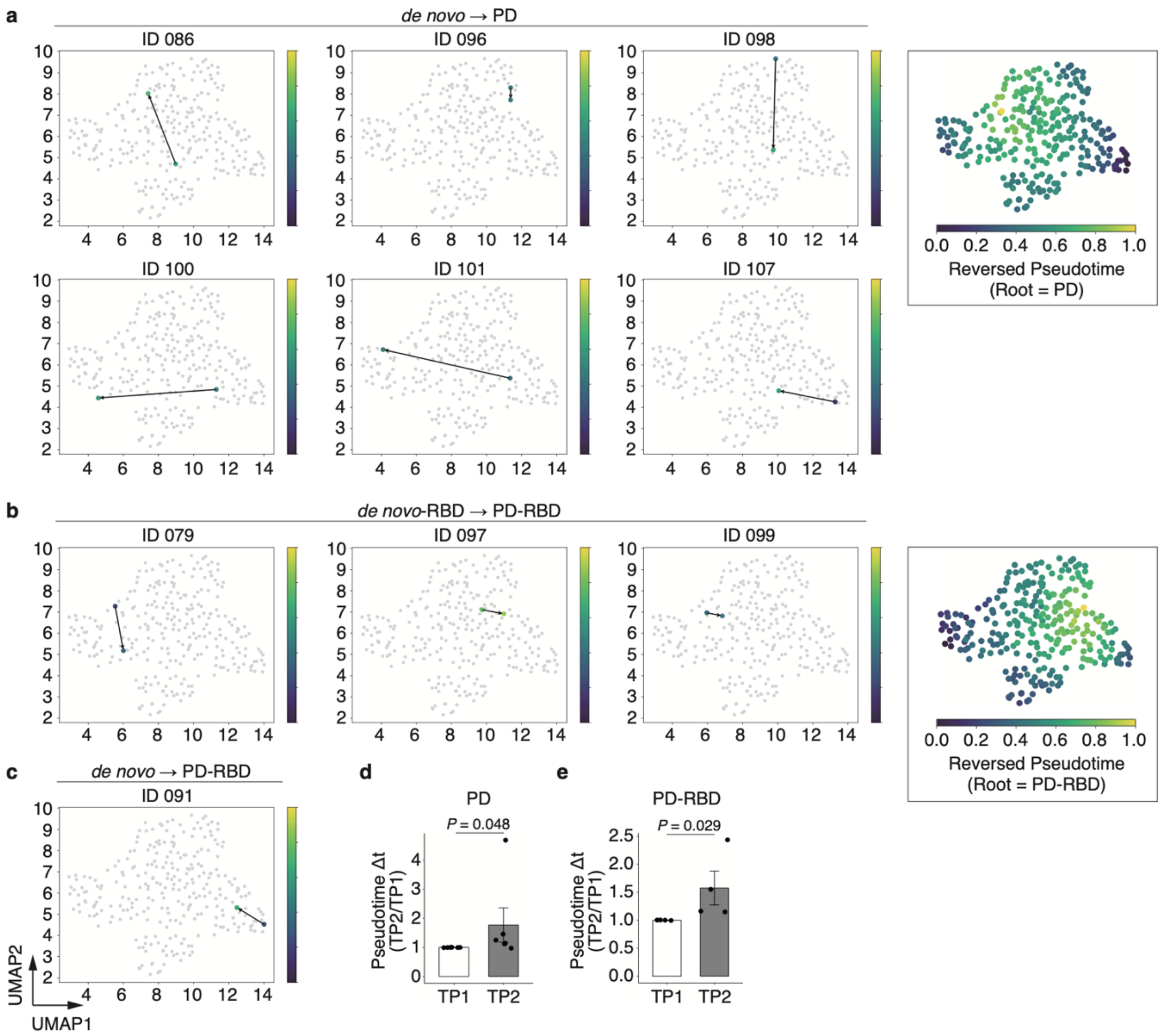


**Supplementary Fig. 4. Trajectory analysis with reversed pseudotime in representative *de novo* and *de novo*-RBD patients.**

**a–c**, Scatter plots showing the trajectory of the longitudinal analysis (after treatment with anti-PD drugs) in *de novo* patients on the manifold with reversed pseudotime. The gray dots represent visualization of the microbiome compositional manifold, and colored dots indicate the locations of each sample of patients transitioned from *de novo* to PD (**a**), *de novo*-RBD to PD-RBD (**b**), or *de novo* to PD-RBD (**c**). Black arrows indicate the direction of the sampling time. **d**,**e**, Graphs showing changes (Δt) in each reversed pseudotime rooted at PD ($n$=6; **d**) or PD-RBD ($n$=4; **e**) between two sampling times (TP1 and TP2). Dots indicate each individual. Statistical significance was determined using the Wilcoxon rank-sum test ($P$<0.05).

*de novo*, *de novo* patients with PD; PD, Parkinson's disease; RBD, rapid-eye-movement sleep behavior disorder; TP, time point.

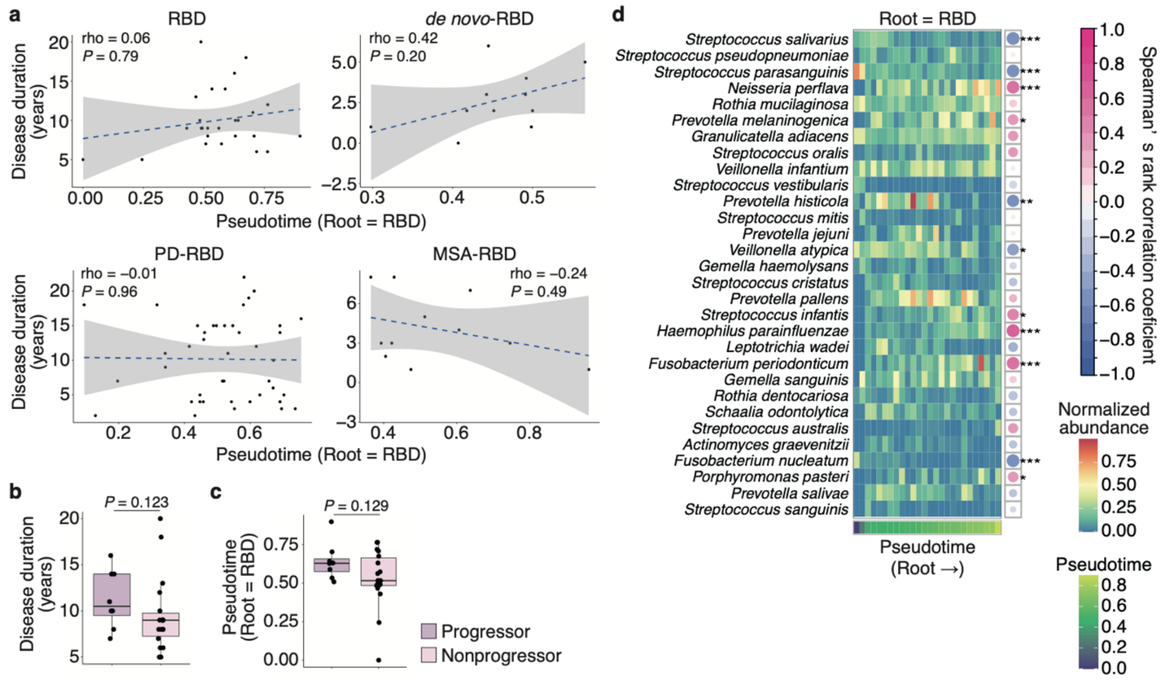


**Supplementary Fig. 5. Shifted microbial abundance with pseudotime in patients with RBD.**

**a**, Graphs showing the correlation between each disease duration and pseudotime (RBD as the root). Statistical significance was determined using Spearman's correlation coefficient ($P$<0.05). **b**,**c**, Graphs showing the disease duration of RBD (**b**) and pseudotime rooted at RBD (**c**) between patients who have developed synucleinopathies (progressors) or still have RBD (nonprogressors). The box plots present the median and 25th and 75th percentiles. Dots indicate each individual. Statistical significance was determined using the Wilcoxon rank-sum test ($P$<0.05). **d**, Heatmap showing the normalized abundance of the top 30 species (a relative mean abundance of >0.1%) along with pseudotime (RBD as the root). The right panel represents Spearman's correlation coefficient between normalized abundance and pseudotime. Statistical significance was determined using Spearman's correlation coefficient ($P$<0.05). *$P$<0.05; **$P$<0.01; ***$P$<0.005.

*de novo*, *de novo* patients with PD; MSA, multiple system atrophy; PD, Parkinson's disease; RBD, rapid-eye-movement sleep behavior disorder.

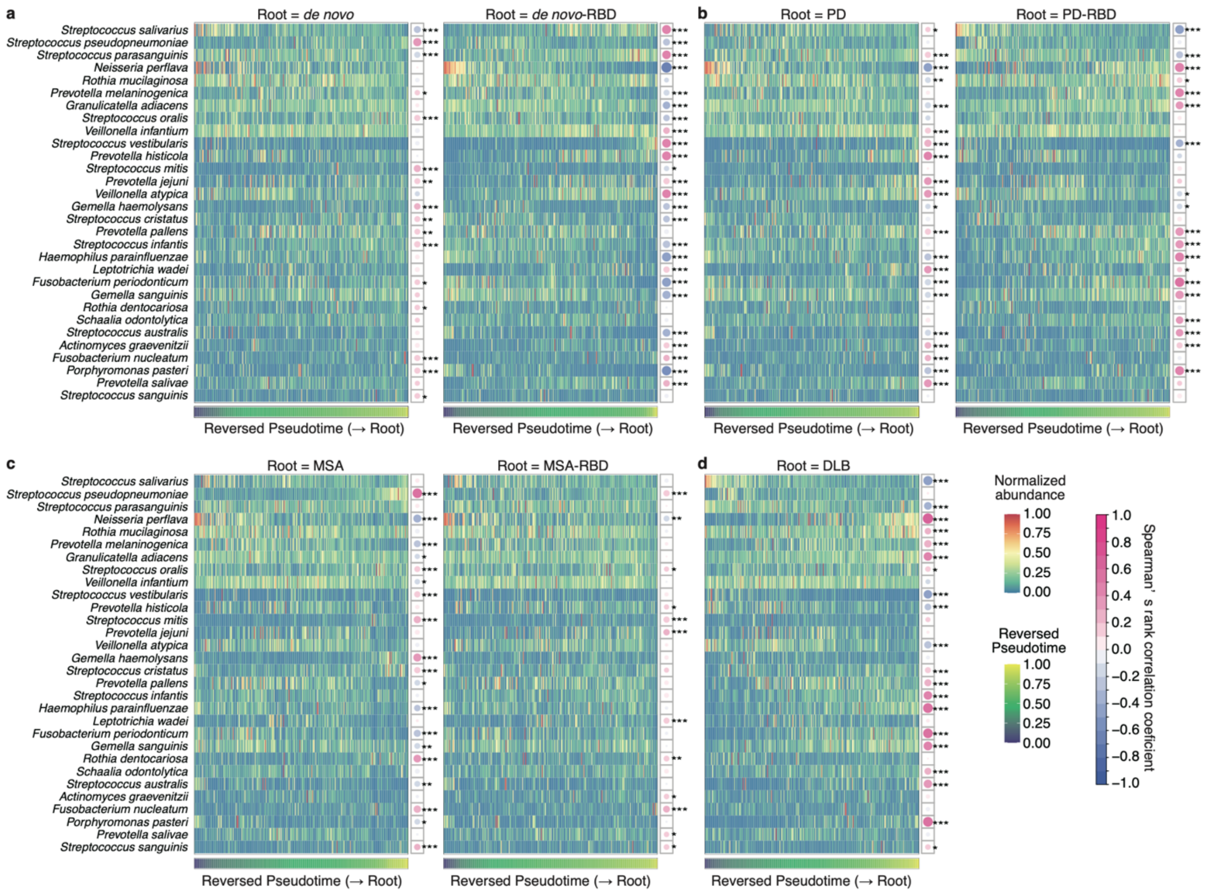


**Supplementary Fig. 6. Microbial abundance altered with reversed pseudotime in patients with synucleinopathies.**

**a–d**, Heatmaps showing the normalized abundance of the top 30 species (a relative mean abundance of >0.1%) along with reversed pseudotime rooted at *de novo*, *de novo*-RBD (**a**), PD, PD-RBD (**b**), MSA, MSA-RBD (**c**), and DLB (**d**). Each right panel represents Spearman's correlation coefficient between normalized abundance and reversed pseudotime. Statistical significance was determined using Spearman's correlation coefficient ($P$<0.05). *$P$<0.05; **$P$<0.01; ***$P$<0.005.

*de novo*, *de novo* patients with PD; DLB, dementia with Lewy bodies; MSA, multiple system atrophy; PD, Parkinson's disease; RBD, rapid-eye-movement sleep behavior disorder.

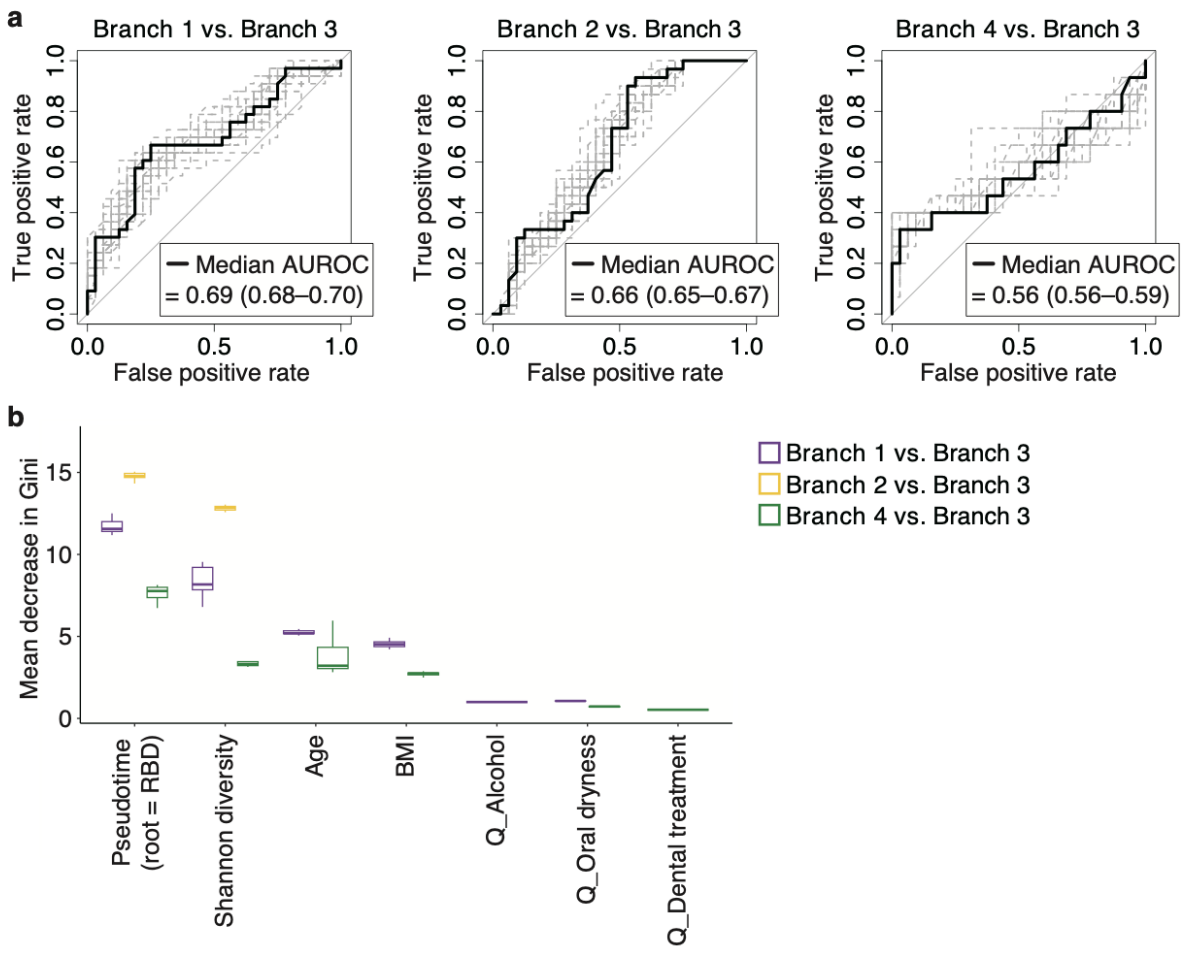


**Supplementary Fig. 7. Host factors in branch 3 with a short pseudotime from prodromal to early PD stages.**

**a**, Graphs showing the ROC curves for classifying branch 3 from the other branches. The distribution of each branch is as follows: branch 1=33, branch 2=30, branch 3=32, branch 4=15. The random forest classifiers take as input pseudotime (RBD as the root), Shannon diversity score, and host factors (age, sex, BMI, questionnaire responses [dental treatment, toothbrushing, denture, oral dryness, smoking, and alcohol consumption]). For calculating median AUROC and 95% CIs (shown in the figure), 25 iterations of independent training were performed using different random seeds. Dashed gray lines represented the results from 25 iterations, and bold lines indicated the median. **b**, The mean decrease in the Gini index, representing variable importance, was obtained from 25 independent iterations.

AUROC, area under the receiver operating characteristic curve; BMI, body mass index; Q, questionnaire; RBD, rapid-eye-movement sleep behavior disorder.

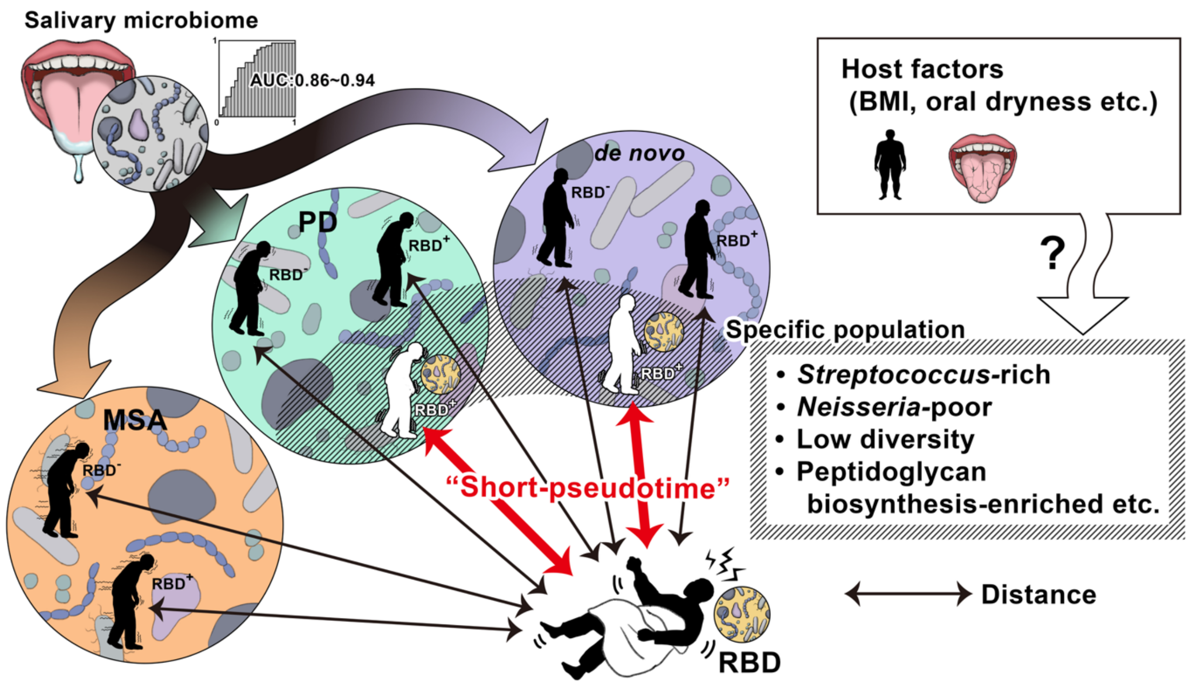


**Supplementary Fig. 8. Graphical summary.**

Schematic illustration of the main results that the salivary microbiome profiles allow for stratification of synucleinopathies based on the high AUC value. Notably, the microbiome composition in patients with RBD is similar to that in patients with synucleinopathies presenting RBD symptoms, rather than in those without RBD symptoms. A short pseudotime from RBD to the early PD stage was observed in a specific population characterized by *Streptococcus*-rich, *Neisseria*-poor, and low-diversity profiles. The KEGG enrichment analyses revealed metabolic pathways (e.g., peptidoglycan biosynthesis) enriched in the short pseudotime population.

AUC, area under the curve; BMI, body mass index; *de novo*, *de novo* patients with PD; KEGG, Kyoto Encyclopedia of Genes and Genomes; MSA, multiple system atrophy; PD, Parkinson's disease; RBD, rapid-eye-movement sleep behavior disorder.
